# Gene regulatory network inference from single-cell data using optimal transport

**DOI:** 10.1101/2024.05.24.595731

**Authors:** François Lamoline, Isabel Haasler, Johan Karlsson, Jorge Gonçalves, Atte Aalto

## Abstract

Modelling gene expression is a central problem in systems biology. Single-cell technologies have revolutionised the field by enabling sequencing at the resolution of individual cells. This results in a much richer data compared to what is obtained by bulk technologies, offering new possibilities and challenges for gene regulatory network inference. Here we introduce GRIT — a method to fit a differential equation model and to infer gene regulatory networks from single-cell data using the theory of optimal transport. The idea consists in tracking the evolution of the cell distribution over time and finding the system that minimises the transport cost between consecutive time points.

## 1 Introduction

Understanding cellular functions and exploring the regulatory relationships between genes is a central problem in systems biology and medicine. This is key to reveal the molecular mechanisms of specific diseases and to find treatments eventually. Emerging single-cell techniques now allow researchers to perform sequencing at the resolution of individual cells for a large number of cells at a time. However, the measurements are destructive, which prevents observing a cell over time. Instead, the data consist of population snapshots at different times.

Inference of gene regulatory networks (GRNs) from bulk time series data has been a longstanding problem in systems biology [44, 2, 31]. Inference of dynamical models from time series data is also a common problem in many fields of engineering. However, compared to most engineering applications, in molecular biology, data collection is expensive and laborious. Therefore, the main challenge in GRN inference from time series data is to deal with the small amount of data, in particular, given the high dimension of the problem. In the highly simplified framework of discrete-time linear systems with full state measurements, it is well known that to infer an *n*-dimensional system, at least *n* + 1 time points are needed. This requirement is never satisfied with bulk transcriptomics data, unless the set of genes to include is heavily restricted. A common workaround is to impose sparsity constraints to the inferred models.

Single-cell data, in contrast, reveal the time evolution of the full distribution of the cell population. In [3], it is shown that three time points of such population snapshot data are sufficient for unique identifiability of a discrete-time linear system, regardless of the system dimension. Despite the overly simple model class, this result gives hope that single cell data can accelerate transcriptomics research. Methods for GRN inference from single-cell data have been proposed, based on, for example, information-theoretic considerations [16], regression [31, 47], co-expression [62], and dynamical models [45, 1, 51, 9, 65]. In a benchmarking study [55], regression-based methods seemed to be the most consistent, and were the best performers in the inference task from real scRNA-Seq data. Regression-based methods — even though demonstrated to have good performance — are heuristic, and not rooted in physical considerations. Moreover, they do not directly extend to other tasks related to GRN inference, such as finding targets of perturbations in the network.

This work uses the concept of optimal transport (OT), a theory that provides a natural way to compare two given distributions. The original formulation of the problem, due to Gaspard Monge in 1781 [48], asks to find a transport mapping from one distribution to the other, such that a given total transportation cost is minimised. While Monge introduced the problem discussing a very concrete problem of moving earth from an initial configuration to a target configuration, more recently optimal transport has found numerous applications ranging from machine learning [38] to imaging, probability theory [58] as well as systems theory [17, 18]. With the advent of single-cell data, OT has also found several applications in systems biology. In [28, 12], OT is used to infer cell dynamics driven by a gradient flow governed by a potential function (analogous to the famous Waddington landscape). In [60], OT is used to find couplings between consecutive time points to identify single cell trajectories in the form of ancestor and descendant distributions. In [29], OT is used to formulate a metric for cell-cell similarity, interpreting each cell’s feature profile as a probability distribution, and defining a cost of transport between features. OT is used in [30] on paired single-cell multiomics datasets as a loss function in a matrix factorisation approach. In [13], OT is used to couple a perturbed cell group with a control group to predict perturbation responses. In [14], OT is used to align single-cell data with spatial transcriptomics data using a few marker genes, and further, to match ligand and receptor distributions to study cell-cell communication (developed further in [15]).

This work introduces GRIT (Gene Regulation Inference by Transport theory), a method based on fitting a linear differential equation model to the observed data using OT. More precisely, the method works by propagating cells measured at a certain time *T*_*k*_ through a candidate model, and calculating the transport cost between the propagated population and the cell population measured at the next time point *T*_*k*+1_. The goal is to determine the model that minimises the optimal transport cost. GRIT naturally extends to the aforementioned additional challenges, namely inference of perturbation targets and mutation effects. Moreover, we prove a consistency result for the method, stating that if the data were generated from a linear discrete-time system, then the true system is the unique global minimiser of the defined transport cost. GRIT is first validated using synthetic data to study the effect of various details on its performance. In particular, we demonstrate GRIT’s ability to infer perturbation targets and mutation effects. GRIT is then applied on the BEELINE benchmarking pipeline [55] to compare it with state-of-the-art methods. GRIT demonstrates robust and good performance, in particular outperforming state-of-the-art methods when applied to simulated data. Finally, GRIT is applied on two real datasets [50, 68], generated with the aim to study the effect of different mutations (*LRRK2*-G2019S and *PINK1*-I368N) associated with Parkinson’s disease on neuron differentiation. GRIT identified perturbation targets corresponding to the mutations as well as enriched pathways and we compared the findings with existing literature.

## 2 Results

### 2.1 GRIT: Gene regulation inference by transport theory

This section introduces GRIT, the method for gene regulatory network inference from single cell time course data. A more detailed description, including a more extensive background on optimal transport theory and rationale for some modelling choices can be found in Appendix A.

The optimal transport (OT) theory is a natural tool for finding a coupling between two probability distributions in the continuous case or two point clouds in the discrete case, and measuring the distance between them. Say 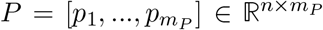 is a matrix containing the points of one point cloud consisting of *m*_*P*_ points and 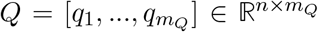 another point cloud consisting of *m*_*Q*_ points. To define the discrete OT problem, a *transportation cost* needs to be defined between each pair of points (*p*_*i*_, *q*_*j*_). In this work, the cost is defined as the squared distance between *p*_*i*_ and *q*_*j*_. The discrete OT problem (with entropy regularisation) is

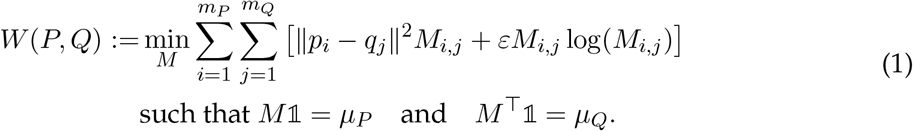

where 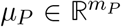 and 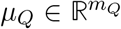 are the weights of the samples in the respective distributions. The *transport plan* 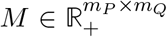 gives a coupling between the points in the two sets that minimises the total cost of transportation. The minimal value is used as a measure of the quality of fit between *P* and *Q*. The entropy regularisation is fairly standard in OT problems with several functions [53, Chapter 4]. For example, it enables efficient numerical solution by the so-called Sinkhorn iterations [22], but in our approach it also accounts for noise in the cell dynamics.

Cells are modelled as individuals evolving in the gene expression space. The gene expression levels of *n* genes form the state vector *x* of a cell. It is assumed to be governed by the stochastic linear differential equation

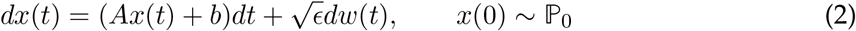

where *A* is a sparse matrix, *b* is a constant load, *w* is a standard Brownian motion, and *ε >* 0 is the noise intensity. Biologically, *A* contains the regulatory parameters of the transcription factors (TFs). As mentioned above, the entropy regularisation in (1) is related to the noise in cell dynamics, and it is not an accident that we use the same symbol *ε* for both noise intensity in (2) and regularisation strength in (1).

As summarised in Figure 1, the optimal transport cost is used to evaluate the performance of a dynamical system (2) in fitting observed data. To this end, the distribution measured at time *T*_*k−*1_ is first propagated through system (2). Then, this propagated distribution is compared with the population measured at *T*_*k*_. To obtain a mathematically tractable problem, the propagation is done using a first-order (Euler) discretisation of (2). If *Y*_*k−*1_ is the expression matrix for the population measured at time *T*_*k−*1_, the propagated matrix is given by (*I* + Δ*T*_*k*_*A*)*Y*_*k−*1_ + Δ*T*_*k*_*b*. The cost function for evaluating a model (*A, b*) is then obtained by combining all time points:

**Figure 1:**
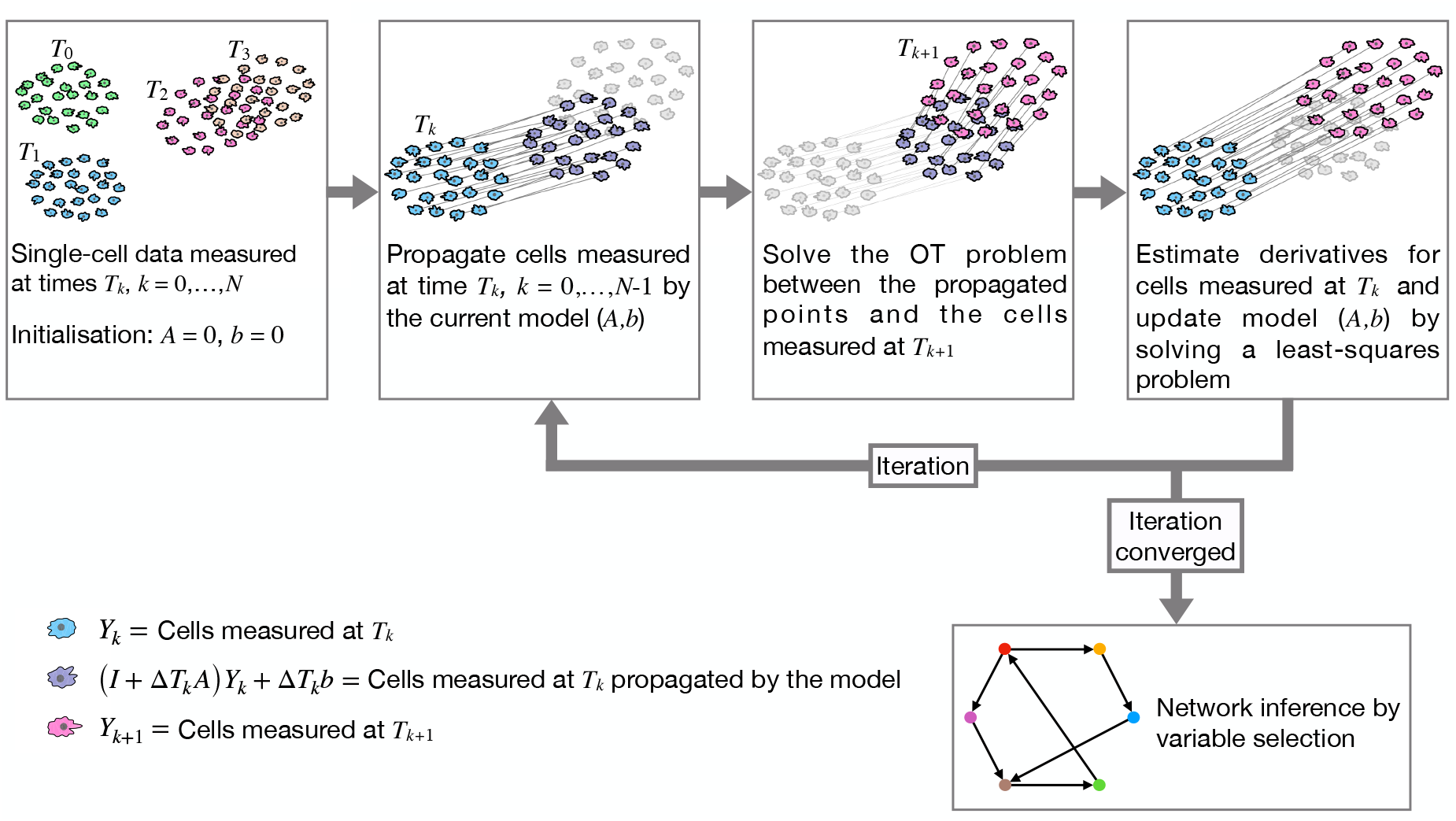
Method pipeline: The model identification iteration is run until convergence. The results are then used in a separate variable selection step to obtain confidence scores for GRN link existence.

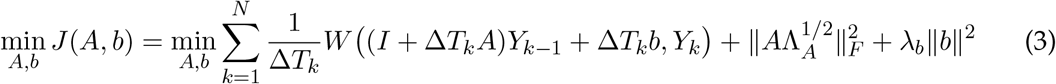

where the diagonal matrix Λ_*A*_ and *λ*_*b*_ are regularisation parameters. The weight 1*/*Δ*T*_*k*_ in the sum is related to the effect of noise *dw* in (2). Note that the calculation of the transport cost *W* involves the solution of a minimisation problem (1). The combined optimisation problem for the system variables *A, b*, and the transport plans *M*_*k*_ for *k* = 1, …, *N* is solved by a coordinate-descent type algorithm, illustrated in Figure 1. When *A* and *b* are fixed, the minimisation problem is a standard OT problem that is solved by Sinkhorn iterations. When, in turn, the transport plans are fixed, the minimisation problem is a quadratic problem with respect to (*A, b*). The full problem is solved by alternately minimising with respect to the two variables.

Our main objective is to use the optimal transport cost as a goodness-of-fit measure for identifying the dynamical system *A, b*. However, the transport plans *M*_*k*_ that are obtained for each time transition *T*_*k−*1_ *→ T*_*k*_ are interesting on their own. They can be used to identify ancestor and descendant cells as is done in the Waddington-OT approach [60]. The transport plan *M*_*k*_ can also be used to define a target point at time *T*_*k*_ for each cell measured at time *T*_*k−*1_. With matrix notation, the target points for all cells in the matrix *Y*_*k−*1_ are given by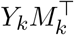 . These target points can then be used to estimate derivatives for the cells (corresponding to RNA velocity [39]) by a difference quotient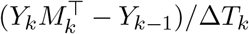 . As shown in Supplementary note 1, the quadratic problem for (*A, b*) can be reduced to a regression problem involving these derivative estimates.

The use of a first-order discretisation effectively renders the method a discrete-time scheme.

If, indeed, the data originate from a discrete-time system, then the following result holds.

**Box 1. Consistency result**

Assume that the data are produced by a discrete-time system

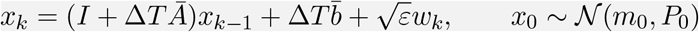

and data on at least three time points *k*Δ*T* with *k* = 0, 1, …, *N* is measured. With the number of cells measured on each time point tending to infinity, the true system 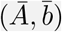 is the unique global minimiser of the cost function (3).

The precise statement of this result and its proof can be found in Supplementary note 2. The proof relies on recent results on entropy-regularised optimal transport [32, 46] and on an identifiability result for uniqueness [3]. Interestingly, as already mentioned, the entropy regularisation acts as a noise deconvolution accounting for the noise in the stochastic dynamics. This idea of entropy regularisation as noise deconvolution holds more generally and it has been explored in [58].

The dynamical system parameterised by the matrix *A* and the vector *b* already describe the regulatory structures between genes. In principle, the higher the absolute magnitude of an element in the *A* matrix, the stronger is the corresponding regulation. However, the matrix element magnitudes are sensitive to gene expression level scales and they are not necessarily the best measure of the confidence on the existence of a particular regulatory effect. Therefore once the model identification algorithm has converged, the results are used in a separate variable selection step, whose results are interpreted as confidence scores for the existence of a regulatory link. The additional variable selection step is based on a greedy forward-backward sweep (inspired by [70] to which we also refer for details on the greedy approach) applied on the regression problem obtained from (3) by keeping the transport plans *M*_*k*_ fixed. To initialise the backward greedy algorithm, the initial set of regressors is selected based on a combination of gene-gene correlations and a forward greedy algorithm, that is, at every step adding to the active regressor set the regressor yielding the highest decrease in the cost function. The backward greedy algorithm is then carried out, that is, at every step removing the regressor that yields the smallest increase to the cost function. The confidence scores for the existence of links are based on cost function increments during the backward sweep.

In some single-cell datasets, the population splits into separate subgroups over time, for example, differentiating into different cell types. In such cases, it is advised to first identify branching dynamics and give branch labels as an input to GRIT. The handling of branching dynamics is described in Section A.5. Branch labels can be obtained using, for example, Slingshot [64] or any other suitable pseudotime inference method, or alternatively, using the GRIT branch labeling scheme described below in Sections 2.7 and A.8.

### 2.2 Performance on linear discrete-time dynamics

As stated in the consistency result in Box 1, GRIT is essentially a system identification tool for linear discrete-time systems from population snapshot data. To investigate the performance of the method in this task, it is here applied on data generated from a 10-dimensional linear discrete-time system (see Supplementary note 3 for details on the data generation). The consistency result holds when the number of cells measured at each time point tends to infinity. However, it should be noted that the consistency result only guarantees that the true system is the unique global minimiser of the cost function (3), but it does not guarantee convergence to this minimum. To study convergence properties, GRIT was applied to data with either three, six, or twelve time points, with the number of cells per time point being 1000, 2000, 3000, 5000, 7500, or 10000. In this experiment, the method was tuned to correspond precisely to the assumptions of the theorem, that is, we set Λ_*A*_ = *λ*_*b*_ = 0, and the entropy regularisation parameter *ε* was set to the noise intensity used in the data generation. The results are shown in Figure 2a in terms of squared Frobenius norm between the estimated and true (*A, b*). The estimated system appears to converge towards the true system such that 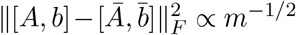 where *m* is the number of cells per timepoint. Incidentally, this is precisely the convergence rate (in matrix norm) of the empirical covariance matrix [66]. The computation time is polynomially increasing (*∝m*^2.42^) with the number of cells (Supplementary figure 7).

**Figure 2:**
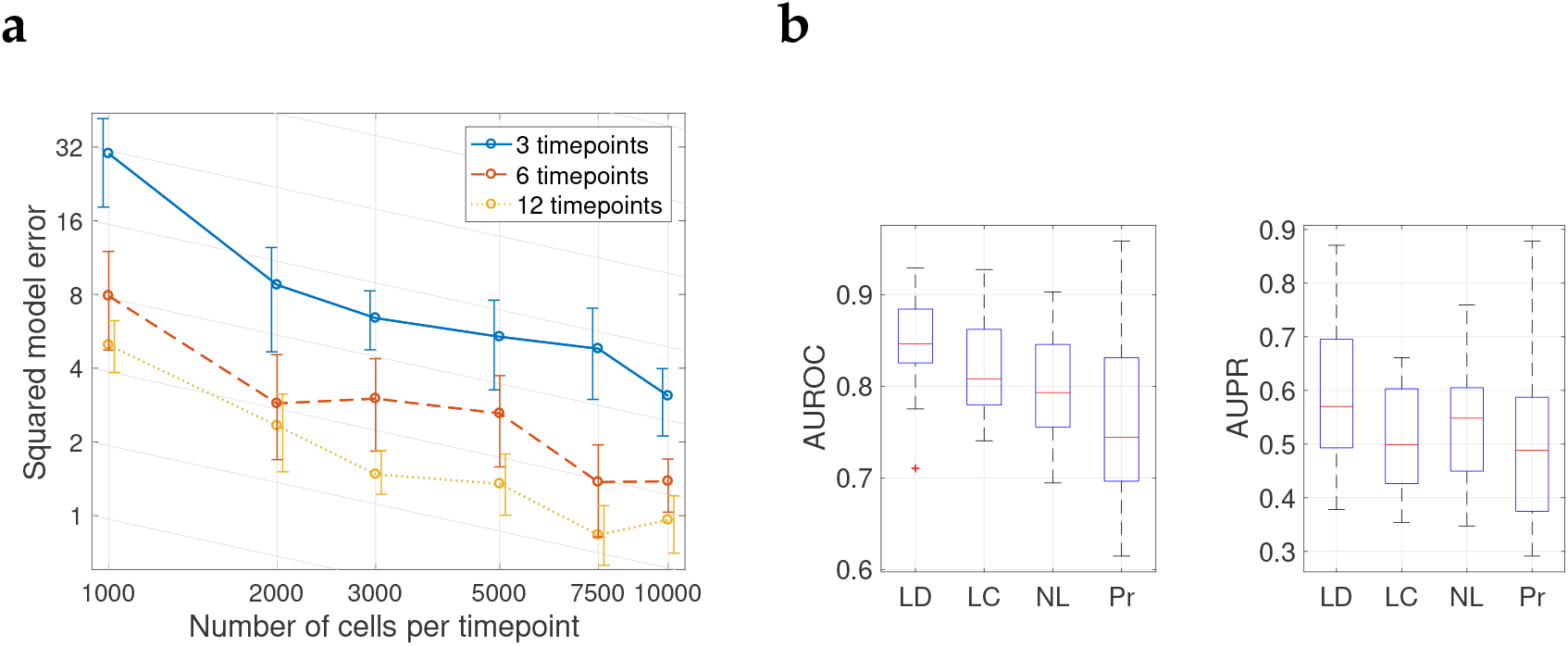
**a:** The squared model error 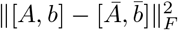 with data simulated from a linear discrete-time system with varying number of timepoints and number of cells per timepoint. The plot is in logarithmic scale and it shows the mean and 80th percentiles obtained from five replicates. The sloped gridlines correspond to a decay *m*^*−*1*/*2^. **b:** The AUROC and AUPR scores from 20 replicates with data simulated from linear discrete-time system (LD), linear continuous-time system (LC), nonlinear system (NL), or system with non-observed protein concentrations (Pr).

### 2.3 Effect of continuous time, nonlinear dynamics, and hidden states

Obviously, the framework of linear discrete-time dynamics is a crude simplification of underlying biological processes. The goal now is to go step-by-step towards a more realistic setup and study how the performance of GRIT changes. First, continuous-time simulations are done instead of discrete-time. Then, the linear dynamics are replaced by nonlinear dynamics including transcription saturation by Michaelis–Menten kinetics as well as more realistic inhibition action. Moreover, the dynamics are corrupted by state-dependent Langevin noise, which is a more realistic noise model than constant-intensity white noise used with the linear system [23]. Finally, hidden states mimicking protein concentrations are introduced, one protein corresponding to each gene with linear dynamics for the protein translation and degradation. Further details of these systems can be found in Supplementary note 3.

Results on 20 replicates are shown in Figure 2b. In this experiment, six time points have been simulated with measurement times equally spaced with 0.4 time units in between. There is a clear drop in performance when moving from discrete- to continuous-time systems. This is somewhat expected when moving away from the precise model class for which the method should work perfectly, up to noise and incomplete information due to finite sample sizes. Interestingly, there is no significant decrease when moving from linear to nonlinear dynamics.

The performance drops again when hidden states are introduced. Moreover, the variability in performance is quite high in this case, which is most likely due to the randomisation of the parameters in the protein dynamics. Slower protein dynamics cause delays that might blur the observability of regulatory interactions between genes.

This experiment was also used to test the effect of using single-cell data instead of corresponding bulk data. To this end, GRIT was applied on data where the expression vector of each cell was replaced by the average expression level of the corresponding time point. This corresponds to running GRIT’s network inference step on bulk data. The performance is far worse than with single-cell data (Supplementary figure 1a). In addition, the variable selection step for network inference was validated. Using the absolute values of the *A*-matrix entries as confidences for link existence, the results are reasonable but still inferior to the GRIT results (Supplementary figure 1b).

One should not make too far-reaching conclusions based on one example. Indeed, the measurement frequency, nature of the nonlinearities in the system (and transcription saturation level), and time scales of protein dynamics all have a considerable effect on the quality of the inference results.

### 2.4. Perturbation target inference

A standout feature of a differential equation based method is the ability to deal with perturbations in a straightforward manner. Given data on a control experiment and a perturbation experiment, GRIT can infer targets of the perturbation (for example, a drug). Perturbation target inference by GRIT is explained in Section A.6. Essentially, an additional vector *b* is added to (2) that is active only in the perturbation dataset.

To validate the inference of targets of an external perturbation, data were generated from the modified nonlinear system presented in Supplementary note 3. In the modification, the basal transcription rates of different genes were modulated. Six cases were generated where either one, two, or three genes were affected by the perturbation with strength of either 30% or 60% increase in the basal transcription rate. Each case consisted of 20 replicates with a control experiment simulated from the model without any perturbations, and a perturbation experiment. The perturbation targets were randomly chosen in each replicate.

GRIT is given the data for both the control and perturbation experiments simultaneously and it is informed that the second experiment has a perturbation. GRIT then outputs an additional vector consisting of confidence values on which genes’ dynamics are likely directly affected by the perturbation. These vectors were concatenated for all replicates into one 10 *×* 20 matrix, that could be compared to a ground truth matrix. The results for the four cases are shown in Figure 3a. To provide a reference to compare with, we also included results obtained by calculating gene-gene correlations separately for the control and perturbation experiments, and summing up the absolute differences of correlations over each row of the correlation matrix. These differences should pinpoint changes in gene dynamics.

**Figure 3:**
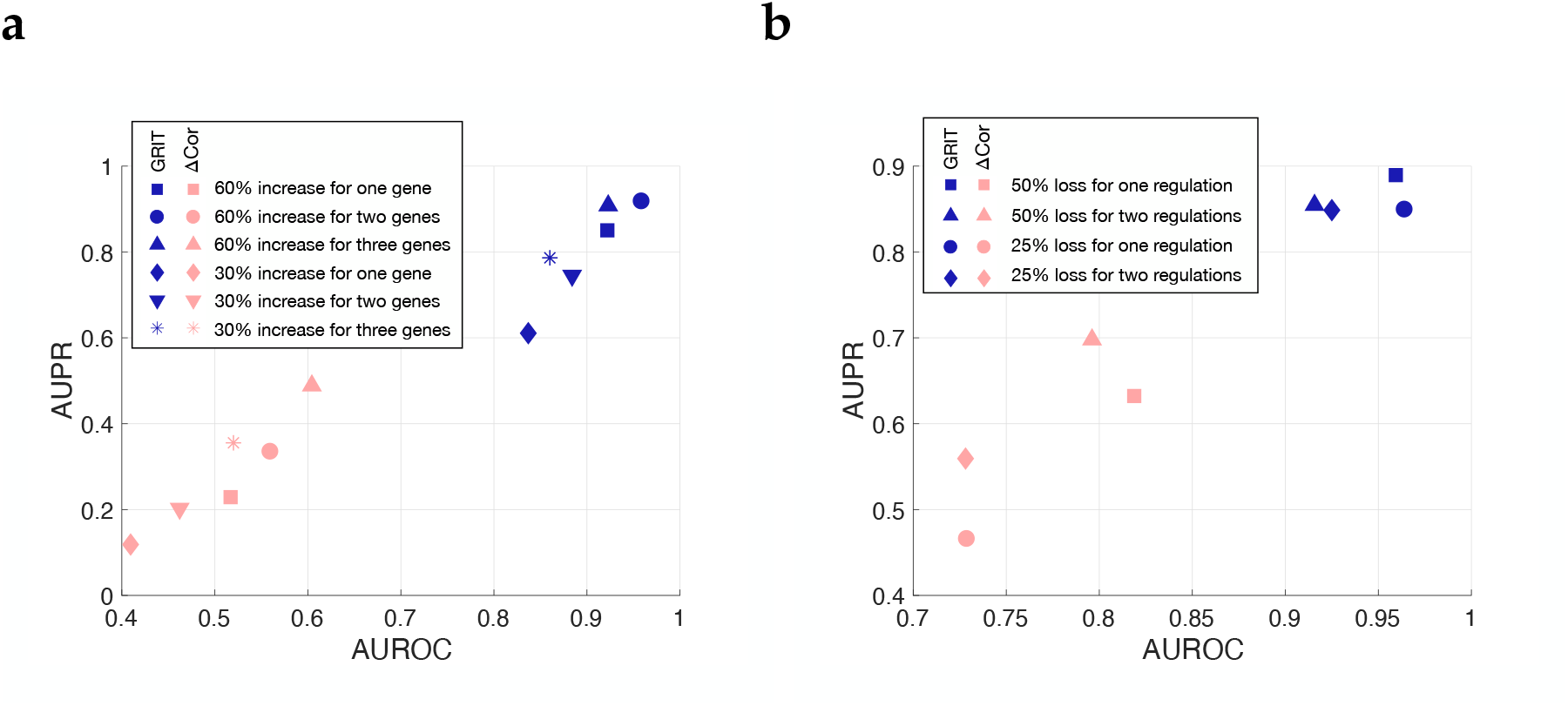
**a:** Results for the inference of perturbation targets by GRIT and by differences in gene-gene correlations in the six different experiments. **b:** Results for the inference of mutation effects by GRIT and by differences in gene-gene correlations in the four different experiments.

### 2.5 Mutation effect inference

GRIT can be used to identify regulations that are altered due to a mutation in a known gene. As with perturbation target inference, a control dataset and a mutation dataset are needed, that should otherwise correspond to the same experimental conditions, and preferably should be isogenic except for the mutation. Such datasets can be generated by gene editing that allows inflicting targeted mutations into a cell line. The mutation effect inference by GRIT is explained in Section A.7. Essentially, the column in the matrix *A* corresponding to the mutated gene is allowed to differ between the control and mutation datasets.

To validate the inference of affected regulations due to a mutation, data were generated from the modified nonlinear system (details in Supplementary note 3). In the modification, gene 9 directly regulates three other genes. Four cases were generated where either one or two of the regulations from gene 9 were modulated by a coefficient 0.5 or 0.8 corresponding to a 50% or 20% loss of function for the corresponding regulations. Each case consisted of 30 replicates with a control experiment simulated from the model without any perturbations, and a mutation experiment. In the 30 replicates of each case, each gene or each two gene combination was modulated in 10 replicates.

As with the perturbation target inference, GRIT is given both the control and mutation experiments simultaneously and it is informed of the mutation of gene 9 in the second experiment. GRIT outputs an additional vector consisting of confidence values on which genes’ dynamics are likely affected by the mutation. These vectors were concatenated for all replicates into one 10 *×* 30 matrix, that could be compared to a ground truth matrix. The results for the four cases are shown in Figure 3b. As with the perturbation experiment, we also included reference results obtained by calculating gene-gene correlations separately for the control and mutation experiments, and looking at the absolute differences of correlations corresponding to the mutated gene 9 between the two experiments.

### 2.6 Validation by BEELINE pipeline

The BEELINE pipeline [55] was proposed as a systematic pipeline to evaluate GRN inference methods from single-cell data. It consists of three main datasets (detailed in Section B.1), two of which consist of simulated data with known ground truth networks, and one real data with ground truth networks either extracted from STRING database or obtained by ChIP-Seq. Summary results of GRIT on the BEELINE pipeline are shown in Figure 4 (full results can be found in Supplementary figures 2–6 and Supplementary table 1; see also Section B.2 for details on performance evaluation).

**Figure 4:**
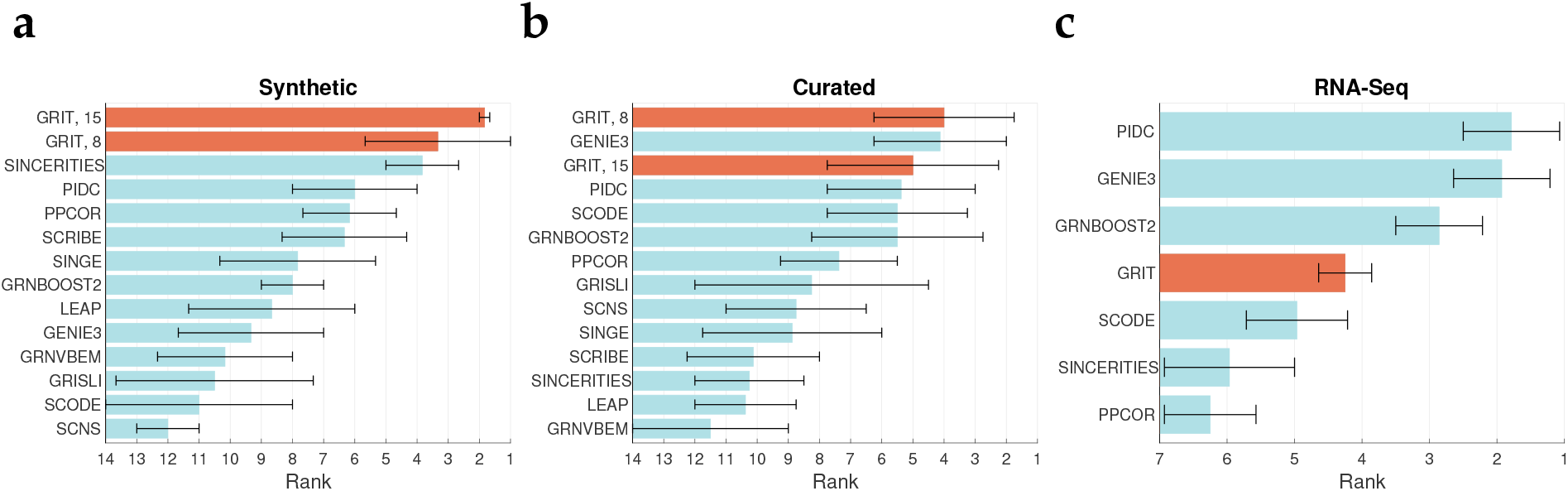
Results summary on the BEELINE benchmarking pipeline for (**a**) the synthetic dataset, (**b**) curated dataset and (**c**) the RNA-Seq data. The bars show the average rank of each method in the different tasks, and the error bars indicate the means of the bottom 50th percentile and the top 50th percentile.

The simulated data in BEELINE are not given in snapshots at different time points, but we have generated time points by ordering the cells by pseudotime (provided along the data) and then dividing the data into eight or fifteen timepoints. These correspond to the results labelled “GRIT, 8” and “GRIT, 15” in Figure 4a–b. Overall, this number of time points did not change much the results (see Supplementary figures 2 and 3), with the exception of the LL system in the synthetic dataset. This system is a cascade of 18 genes that are activated one after the other in a very rapid succession. Eight timepoints is not sufficient to properly capture these fast dynamics resulting in lower performance. A closer inspection of the results revealed that GRIT in this case was inferring many indirect regulations with higher confidence than the true direct regulations (that is, giving high confidence to a regulation *A → C* when the true regulation chain was *A → B → C*). In three of the six systems (BF, BFC, and TF), GRIT outperforms other methods with a large margin. Interestingly, these are the systems with branching dynamics, which GRIT can clearly handle well despite its reliance on linear dynamics. It should be noted that branch information should be given to the method to improve its performance. With the LI and CY systems, GRIT is among the top performers with few other methods that attain more or less perfect score.

In the four inference tasks with the curated dataset, GRIT is the top performer in one task (GSD), among the few top performers in one (HSC), and slightly behind the top performers in two tasks (mCAD and VSC). In the mCAD task, for unknown reasons, very few methods get a score better than random guessing (AUPR-ratio *>* 1). In the data simulated from the VSC model, the data quickly converge to the statistical steady state with dynamical information only in the very beginning. This may favor information theory- and regression-based methods. Re-defining the time scale of the simulation might improve performance for all methods, but particularly for those based on dynamical modelling like GRIT.

Regression-based methods are the best performers in the network inference tasks from real scRNA-Seq data. Curated network databases, like STRING, may have a bias towards correlation- and regression-based methods, since earlier discoveries have often been based on co-expression studies. It is also possible that regression-based methods are more robust against data issues due to limitations of current sequencing technologies. GRIT’s good performance with the cleaner simulated data gives hope of performance improvement as sequencing technology advances. The impact on performance following different transformations is rather small (Supplementary figure 6). The results shown in Figure 4c are obtained with logtransformed data (as provided in [54]).

Following the presentation in [55], the results based on the synthetic data include the cases with 2000 or 5000 cells, and the results based on the curated data include only cases without dropouts. The effects of number of cells and the dropout rate on the performance are illustrated in Supplementary figure 5. Computation times for cases with different dimension and varying number of cells are illustrated in Supplementary figure 7.

**Figure 5:**
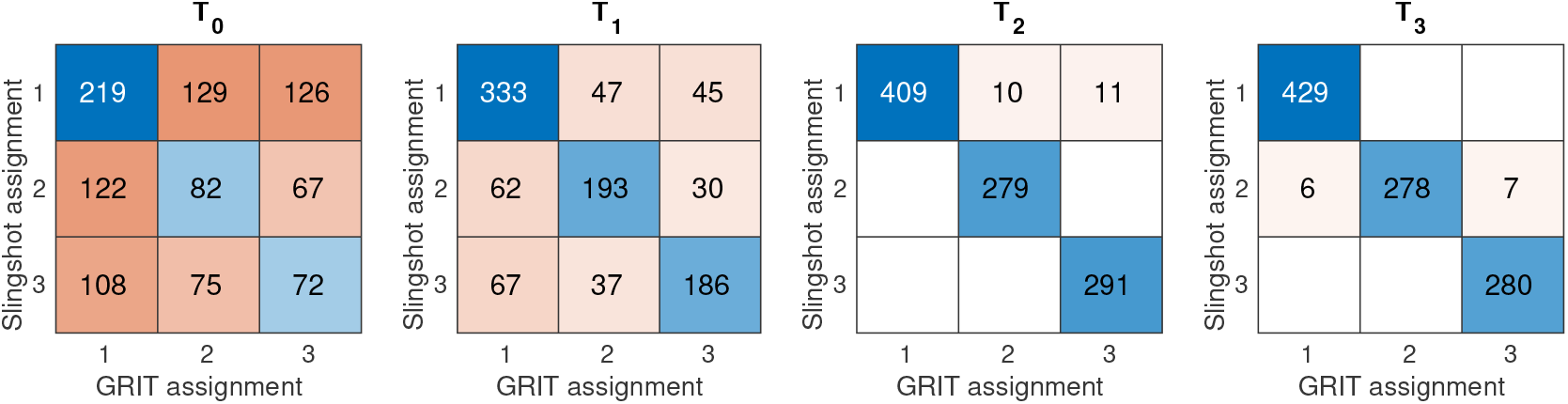
Confusion matrices for branch assignments by GRIT compared to branch assignments of Slingshot at different time points *T*_0_ to *T*_3_. Note that branch assignments from Slingshot at time *T*_4_ were used as cluster labels for GRIT to ensure a meaningful comparison, and therefore a confusion matrix for *T*_4_ is not shown.

### 2.7 Lineage tracing by transport plans

Reaching a complete understanding of a differentiation process or disease progression requires the ability to analyse cell paths. However, due to the destructive nature of the single-cell measurements, cell paths cannot be directly observed. The transport plans *M*_*k*_ can be used for lineage tracing, in particular, for probabilistic identification of ancestor and descendant cells.

In Section A.8, we introduce a scheme that constructs branch labels by tracing the ancestor lineage of cell clusters defined on the final measured time point. More specifically, GRIT is first applied without the branch labels, then cells measured on the final time point are clustered, and finally branch labels are constructed by identifying cells at earlier time points whose descendants belong to the identified clusters of the final time point. This branch labeling scheme is here illustrated on the simulated data of the system producing trifurcating behavior in the BEELINE benchmark pipeline. In this dataset, the initial population is unimodally distributed, and over time the population splits into three subpopulations. Branch indicators obtained by Slingshot are provided with the data (three branches corresponding to the three subpopulations). The data are divided into five time points. The GRIT branch labeling scheme was run by using the provided Slingshot branch labels for the final time point *T*_4_ as the cluster labels. The branch labels from GRIT for time points *T*_*k*_, *k* = 0, 1, 2, 3, can then be compared to the Slingshot branch labels. The confusion matrices for different time points are shown in Figure 5. It can be noted that the identified ancestors respect the branch assignment by Slingshot very well, except for the time point *T*_0_. This, however, is not surprising since the initial distribution is unimodal, and therefore the branch assignments are arbitrary. One should also keep in mind that Slingshot branch assignments are by no means a “ground truth”, but the result simply demonstrates a high level of agreement between the two methods.

### 2.8 Perturbation target identification for *PINK1* and *LRRK2* mutations

In the analysis of the *PINK1* and *LRRK2* results, the focus is put on perturbation target inference, that is, discovery of genes whose dynamics have been directly affected by the mutations. Results with 2000 most highly varying genes are shown. With smaller number of included genes tested, the results (not shown) were very similar (in terms of pathway enrichment). Recall that since *LRRK2* itself was hardly expressed in the data, this experiment had to be treated as a perturbation dataset. *PINK1* is better expressed, but it is not among the 2000 most highly varying genes. *PINK1* is nevertheless included in the analysis. GRIT is applied to the *PINK1* dataset using both the perturbation target inference and mutation effect inference approaches. Figure 6 shows the histograms of the scores of each gene for being affected by the perturbation/mutation and lists the most highly scoring genes.

**Figure 6:**
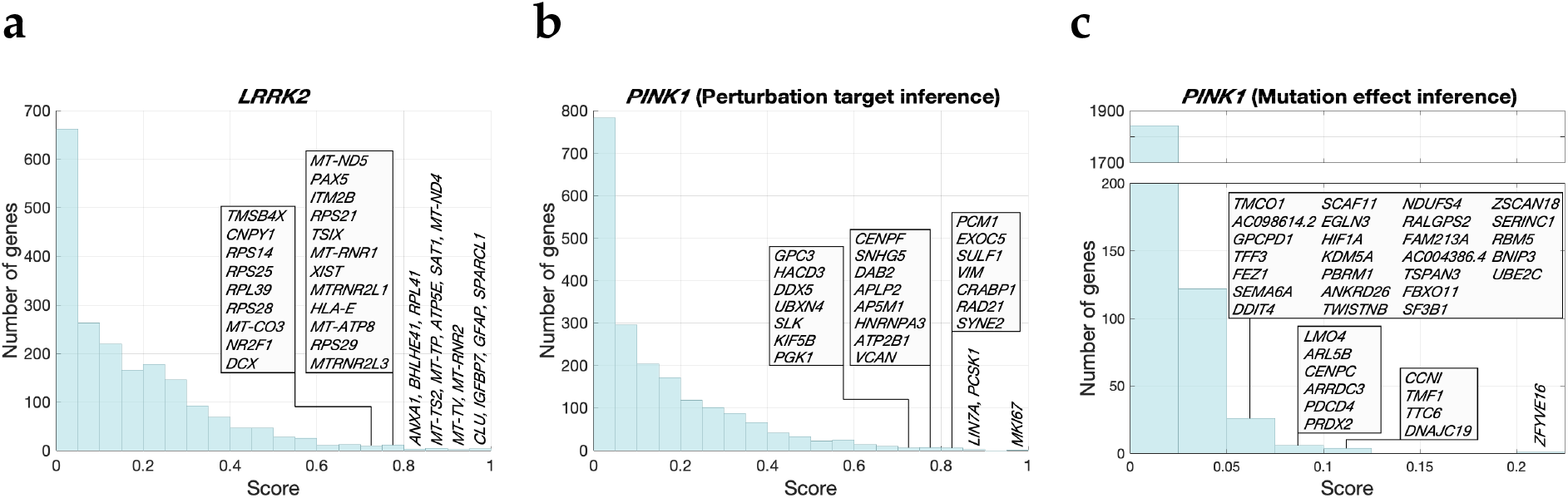
**a:** Histogram of perturbation target scores for the *LRRK2* mutation (**a**) and for the *PINK1* mutation (**b**), and for mutation effect scores for the *PINK1* mutation (**c**). Genes with confidence score above 0.7 are indicated in panels **a** and **b** and above 0.05 in panel **c**.

Of the top-scoring genes in the *LRRK2* case, *SAT1* is PD-associated [40], although without a known direct connection with *LRRK2*. Dysregulation of *ANXA1* by the *LRRK2*-G2019S mutation has been observed previously, although a knockout experiment did not indicate a role in neurodegeneration [57]. *BHLHE41* is a circadian clock gene associated with sleep disorders. Its expression has been found to correlate strongly with the expression of *LRRK2* in [21] where the authors suggest this as a pathway potentially responsible for the narcoleptic-like symptoms in PD. *XIST* has been reported to modulate the *LRRK2* signaling pathway and to accelerate the development of PD [72]. Finally, *NR2F1* is the main gene identified in [68] (where the *LRRK2* dataset originates from) in the mechanism how *LRRK2* influences dopaminergic differentiation.

The top perturbation target candidate in the *PINK1* case with perturbation target inference is *MKI67*, which is a proliferation-related gene inhibited by *PINK1* [33, Figure 4]. Interactions between *PINK1* and *PCM1, VIM, HNRNPA3*, and *DDX5* are reported in [25, Figure 1-1]. Some of the identified genes interact with other well-known PD-associated genes, namely *SNHG5* interacts with *LRRK2* [50], *GPC3* with *DJ-1* (*PARK7*) [50], and *PGK1* with *DJ-1* (*PARK7*) [11].

Of the high-scoring genes in the *PINK1* case with mutation effect inference, *HIF1A* induces mitophagy under hypoxic conditions [42], and a *PINK1* mutation has been reported to affect its transcriptional activity [41]. *GPCPD1* and *BNIP3* are, in turn, regulated by *HIF1A* [42]. A direct interaction between *BNIP3* and *PINK1* has also been described previously and a connection with PD has been established [71]. Interactions between *PINK1* and *PRDX2* and *SF3B1* are reported in [25, Figure 1-1]. *NDUFS4* knock-out induces non-motor symptoms of PD, but no direct connection with *PINK1* is known [20].

To gain overarching insight into the results instead of just looking at the top candidate genes, pathway enrichment analysis was performed on the results with correction for bias due to the selection of genes in the analysis and accounting for the scores given by GRIT (see Section B.3 for details). The mutation effect inference can only detect genes to which a link from *PINK1* has been inferred, and therefore only a handful of genes get a score that is clearly nonzero. Therefore, these results did not yield any significantly enriched pathways (when corrected for multiple hypothesis testing). The results of the analysis for the perturbation target inference results are in Supplementary table 2 for the *LRRK2* case and Supplementary table 3 for the *PINK1* case.

In the *LRRK2* enrichment results, the statistically significant results can be divided in three clusters. The *ribosome* and *COVID-19* terms overlap almost completely (76 genes in common out of 80 genes in the *ribosome* term and 85 genes in the *COVID-19* term). The *cell cycle* term does not have much overlap with any other term, but the remaining 10 terms are highly overlapping with each other. There is a set of 53 genes that belong to at least eight out of ten of these overlapping terms. The KS statistic for this set is 0.379 which is higher than for any of the individual terms. Out of these genes, 23 are in top-400 of GRIT’s results. To zoom in on these genes from the fairly high level KEGG pathways, these genes were fed to g:profiler with GO pathways [8, 5], which gives *oxidoreduction-driven active transmembrane transporter activity* (GO:MF), with term size 69 and it contains 19 of the 23 genes. Other highly significant results are *inner mitochondrial membrane protein complex* (GO:CC) which has term size 162 and it contains all of the 23 genes, *respiratory chain complex* containing 21 of the genes with term size 93, and *mitochondrial respirasome* containing 21 of the genes with term size 96. The *LRRK2*-G2019S mutation has been shown to impair mitochondrial respiration [67]. In the additional results (that are not statistically significant after correction for multiple hypothesis testing), there are several pathways related to DNA repair, namely *Base excision repair, Homologous recombination*, and *Nucleotide excision repair*. The *LRRK2*-G2019S mutation has been shown to cause increased DNA damage [24]. The role of dysfunctional DNA repair pathways in context of PD is also discussed in [24].

In the enrichment results for the *PINK1* case, *serotonergic synapse* is at the top of the results, followed by *dopaminergic synapse* at position 4, *cholinergic synapse* at position 8, and *glutamatergic synapse* at position 32. All of these terms have a high overlap with each other, although the *dopaminergic synapse* term is clearly larger than the other three. Out of the 40 terms in the presented results, 13 have an overlap of *>* 75 % with the *dopaminergic synapse* term. Given the connection to the dopamine system, it is not surprising that these include almost all of the addiction terms (*cocaine, morphine*, and *amphetamine addiction*) on the results. Otherwise, there was no clear clustering of terms like in the *LRRK2* case.

The only pathway term appearing in both *LRRK2* and *PINK1* enrichment results is the *IL17 signaling pathway*, although in neither case it is statistically significant after adjustment for multiple hypothesis testing, mainly due to the small size of the term. However, it is worth to note that in both cases, 50% of the genes are in top-200 of GRIT’s results. The connection of IL-17 (and inflammation in general) and PD has been investigated recently [63, 69].

## 3 Discussion

This work proposed GRIT — a GRN inference method based on optimal transport theory. The inference method consists of simultaneous optimisation of the gene expression model and the transport matrices between gene populations measured at different times. The optimal transport theory is a well-established mathematical framework for comparing particle distributions and it offers a clear interpretation of the optimisation problem in terms of finding most likely particle trajectories from ensemble snapshot observations (see Remark A.1). Moreover, the theoretical tractability of the OT framework allowed us to give a mathematical proof of the method’s consistency — albeit in a very simplified model class of discrete-time linear systems.

GRN inference is not the only utility of GRIT. Indeed, the transport matrices can be used for lineage tracing, that is, identification of potential ancestor and descendant cells, as is done in the Waddington-OT approach [60]. In our approach, the model and transport matrices are inferred simultaneously, and the solution of the OT problem is informed by the inferred model, whereas Waddington-OT is based on solving the OT problem directly between cells measured at different times. In addition, GRIT includes a scheme for identifying targets of perturbations and mutations, which is a task beyond the scope of correlation and information theory -based methods.

GRIT was validated on synthetic and biological scRNA-seq datasets, in particular using the BEELINE benchmarking pipeline. On synthetic data, GRIT outperforms the state-of-the-art methods. With real single-cell RNA-seq data of the BEELINE benchmark, information-theoretic methods seem to perform best. Several factors may contribute to this result. Information-theoretic methods ignore the dynamical nature of the data, and instead concatenate all data together, and then typically try to solve the expression level of a gene as a function of all other genes. While it is obviously not possible to observe gene dynamics with such an approach, potentially these methods benefit from better robustness against sampling problems and low time resolution.

GRIT was applied on datasets studying the effect of *PINK1* and *LRRK2* mutations on the development of dopaminergic neurons. In particular, the goal was to infer perturbation targets corresponding to the mutations, that is, to identify genes whose dynamics are directly affected by the mutation. Some of the top genes identified by GRIT are already known to interact with the mutated genes (*PINK1*/*LRRK2*), and many more are known to be differentially expressed between PD patients and healthy individuals. Moreover, results of the pathway enrichment analysis seem highly reasonable. For example, genes related to mitochondrial respiration and DNA repair are enriched in the top predictions of the *LRRK2* mutation analysis and pathways related to the dopamine system are enriched in the top predictions of the *PINK1* analysis.

Single-cell techniques are not able to capture all of the RNAs from cells. Consequently, genes with low expression may appear as zeros in the data. This phenomenon results in zeroinflated data, which is often considered problematic for analysis [36]. To alleviate this problem, we originally devised a weighting scheme that assigns lower weight to zero values in the data when solving regression problems. While this scheme improved performance with the BEELINE curated datasets, where dropouts are randomly introduced, it does not accurately model the dropout phenomenon. In reality, the dropout probability strongly depends on the expression level of the gene. In fact, with the real RNA-seq data in the BEELINE benchmark, the weighting scheme caused a drop in performance. Finally, we opted to not include the weighting scheme in the published method, and all results in the article are obtained without this scheme. We also experimented with data smoothing by a *k*-nearest neighbour approach, but this did not improve performance either.

Regarding data normalisation and transformation, GRIT does not differentiate between scales of different genes. Hence, it is not essential that all genes are on a similar scale. Log-transformation (with not too extreme scaling) was found to perform well in latent structure discovery [4]. From the point of view of dynamical modelling, it should be noted that the square-root transformation conserves the functional form of linear degradation. That is, if 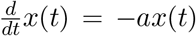, then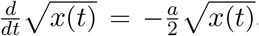 . However, our test with the BEELINE RNA-seq data did not reveal a particularly strong effect of different transformations on the results (Supplementary figure 6).

There are many natural extensions of this framework, such as including growth and death rates of cell populations by considering unbalanced optimal transport, or including cell-to-cell interactions [6] by using the OT formulation and allowing particle interaction [7, 59]. Multiomics integration would provide a thorough comprehension of the regulatory relationships governing cellular mechanisms. For example, fast metabolite dynamics could be modelled by coupling the differential equation model with an algebraic equation based on a quasi-steady state approximation [49]. The so-called RNA-velocity approaches [39, 56] enable estimation of derivatives for the gene expression vectors from the ratio of spliced and unspliced RNA (or, alternatively, using metabolic labeling). While these estimates are rather noisy, RNA-velocity could be integrated into an OT approach to leverage the benefits of both approaches.

## Code availability

Matlab code for the method and data generation are at gitlab.com/uniluxembourg/lcsb/systemscontrol/grit together with a user guide.

## Data availability

The BEELINE benchmark data [54] are available via Zenodo (doi: 10.5281/zenodo.3701939). The PINK1 [50] and LRRK2 [68] datasets are available via GEO with accession code for PINK1: GSE183248 and for LRRK2: GSE128040.

## Acknowledgements

AA and FL are supported by the Luxembourg National Research Fund (FNR) through CORE19/13684479/DynCell.

## A. Method details

### A.1 Model and setup

In this work, R^*k*^ represents the *k*-dimensional real space and ℝ^*k×m*^ the set of matrices of size *k × m*. We denote by 𝟙_*k*_ the all-ones vector of ℝ^*k*^ (though *k* is often omitted if it is clear from the context).

We model the cell population under investigation as a time-varying probability distribution ℙ_*t*_ evolving according to the cellular physiology [27]. Due to the destructiveness of single-cell sequencing technologies, the data are considered as independent samples of distributions 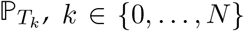 *k ∈ {*0, …, *N }*, where *T*_0_, …, *T*_*N*_ are the measurement times. Denote Δ*T*_*k*_ := *T*_*k*_ *− T*_*k−*1_. The matrices 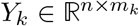 for *k* = 0, …, *N* contain the *m*_*k*_ samples (cells) measured at time *T*_*k*_ where *n* is the number of genes considered.

Cells are modelled as individuals evolving in the gene expression space. Mathematically, the gene expression levels of *n* genes form the state vector *x* ∈ ℝ^*n*^ of a cell. It is assumed to be governed by the stochastic linear differential equation (SDE)

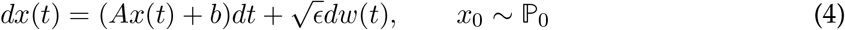

where *A* is a sparse matrix, *b* is a constant load, *w* is a standard Brownian motion, and *ε >* 0 is the noise intensity. The initial state *x*_0_ is assumed to be a realization drawn from a probability distribution ℙ_0_. A comprehensive introduction and background material on stochastic differential equations can be found in [52]. Biologically, *x*(*t*) represents the concentration of mRNA molecules, and *A* contains the regulatory parameters of the transcription factors (TFs). This means that the majority of the elements of the matrix *A* are in fact 0. This knowledge can be used to regularise the problem. Here, we restrict to additive noise in (4), which entails that the random fluctuations are assumed to be independent of the mRNA concentrations. Even though state-dependent noise is more realistic, it can lead to unidentifiable dynamics and overall a more complex problem to solve.

### A.2 Optimal transport

The optimal transport (OT, sometimes called optimal mass transport) theory is a natural tool for finding a coupling between two probability distributions *p* and *q* and measuring the distance between them. It requires the definition of a cost function *c*(*x, y*) (the transportation cost between *x* and *y*) between any samples *x ∼ p* and *y ∼ q*. Optimal transport can be seen as a lifting of the cost function *c*(*x, y*) between samples to a cost function between distributions *p* and *q*. The so-called Kantorovich formulation of the optimal transport problem is to find a joint distribution *μ*(*x, y*) that minimises the total cost of transportation [35] between the distributions *p* and *q*:

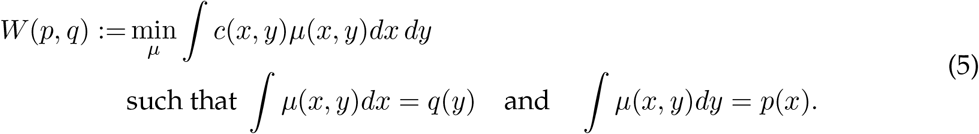

In the case *c*(*x, y*) = *d*(*x, y*)^*r*^ where *d* is a metric, *W* (*p, q*)^1*/r*^ is known as the Wasserstein distance. The solution of the optimal transport problem is the joint distribution *μ*(*x, y*), known as the transport plan — roughly interpreted as the amount of mass transported from *x* to *y* to minimise the total cost. It provides the coupling between the distributions.

In single-cell data, each time point consists of a point cloud of samples from a distribution. For comparison of point clouds, a discrete formulation of the OT problem is used. Say 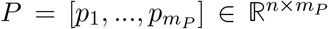 is a matrix containing the points of the first set consisting of *m*_*P*_ points and 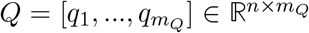 the second set consisting of *m*_*Q*_ points. As in the continuous case, a transportation cost needs to be defined between each pair (*p*_*i*_, *q*_*j*_) and these costs form a cost matrix 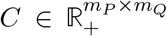. These costs often arise from an underlying cost function, *C*_*i,j*_ = *c*(*p*_*i*_, *q*_*j*_) defined on ℝ ^*n*^ × ℝ ^*n*^. The discrete formulation of the OT problem (with entropy regularisation) is

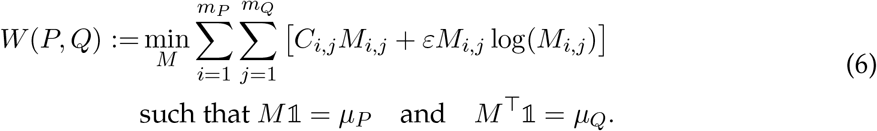

where 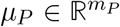 and 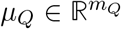 are the weights of the samples in the respective distributions. Note that the discrete formulation arises directly from the continuous case by setting 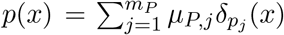 and 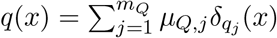 in (5).

The discrete transport plan 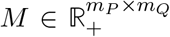 gives a coupling between the points in the two sets. The minimal value is used as a measure of the quality of fit between *P* and *Q*, but due to the entropy regularisation, *W* (*P, P*) ≠ 0, meaning that it is not exactly a distance metric anymore. Below, the term *(entropy-regularised) optimal transport cost* is used for *W* . The entropy regularisation is fairly standard in OT problems and it has several functions [53, Chapter 4]. Firstly, it makes the problem mathematically well-posed (that is, the solution *M* depends continuously on the data *P, Q*). Secondly, it enables the use of the so-called Sinkhorn iterations to solve the problem efficiently [22]. Thirdly, it plays well along with the probabilistic interpretation of the OT problem (see Remark A.1), introducing uncertainty to the OT problem corresponding exactly to the noise process *ε*^1*/*2^*dw* in (4). Note that it is not an accident that we use the same symbol *ε* for the regularisation parameter and the noise intensity. Indeed, it is precisely this last property that underlies the consistency theorem for the method in Section A.3.

As summarised in Figure 1, we use the optimal transport cost to evaluate the performance of a dynamical system (4). The distribution measured at time *T*_*k−*1_ is first propagated through system (4). Then, this propagated distribution is compared with the population measured at *T*_*k*_. To this end, consider a cell trajectory *x*(*t*) satisfying (4) such that *x*(*T*_*k−*1_) = *x*_0_, where *x*_0_ corresponds to a cell measured at time *T*_*k−*1_. Due to linearity of the dynamics, and the Gaussianity of the noise process *w*, at time *T*_*k*_, it holds that *x*(*T*_*k*_) *∼ N* (*m*[*x*_0_], *G*) where

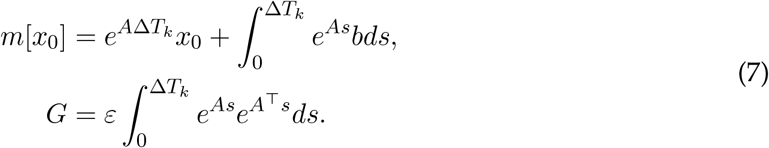

We wish to use the negative log-probability of a cell originating from *x*_0_ (at time *T*_*k−*1_) ending up at *x*_1_ (at time *T*_*k*_) as the transport cost from *x*_0_ to *x*_1_ (see Remark A.1). This is given by

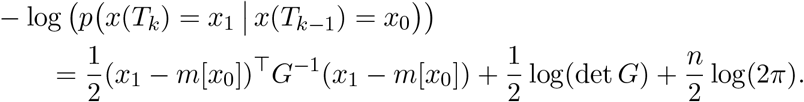

The same cost can be obtained by arguments related to optimal control [17, 18], that is, minimizing the *L*^2^-norm of a perturbation *v* in 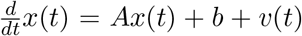 when the initial and final states are fixed to *x*(*T*_*k−*1_) = *x*_0_ and *x*(*T*_*k*_) = *x*_1_. This duality between the linear-quadratic optimal control problem and the Gaussian maximum likelihood problem in the context of OT is also discussed in [26].

Unfortunately, the use of the exponential matrix in the cost function would eventually result in a complex optimisation problem for the estimation of *A* and *b*. To obtain a simpler optimisation problem and to improve robustness of the method, we instead consider a first-order simplification of (7):

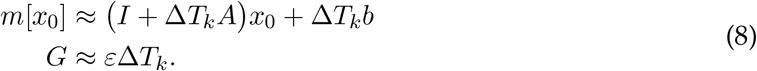

In this simplification, *G* has become a scalar and it no longer depends on *A*. Therefore, the log(det *G*) term (and the *n/*2 log(2*π*) term) can be omitted from the cost. Moreover, the noise intensity *ε* is omitted from the cost as well (the reason becomes apparent in Remark A.1 below), and thus we obtain the transport cost from *x*_0_ to *x*_1_:

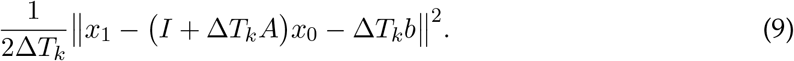

To implement this cost in the transport problem (6), denote by *W* (*P, Q*) the transport cost given by (6), where the underlying cost function is 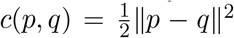. The comparison of data at time *T*_*k−*1_ propagated through the model and the data at time *T*_*k*_ is then done by calculating 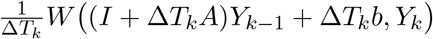.

**Remark A.1**. *As mentioned, the entropy-regularised transport cost is not a distance metric per se, since W* (*P, P*) ≠ 0. *However, the entropy regularisation has an important role in handling noise in the observed distributions. The discrete OT problem* (6) *can be re-written as follows:*

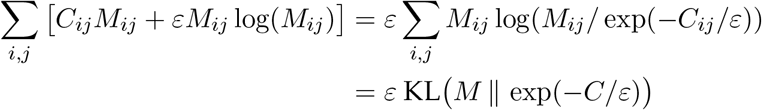

*where* exp(*− C/ε*) *means elementwise exponentation. By definition (modulo the simplification* (8)*), the elements of the matrix* exp(*− C/ε*) *are (proportional to) the a priori probabilities of any cell x*_0_ *from the population measured at time T*_*k−*1_ *ending up to the location of another cell x*_1_ *from the population measured at time T*_*k*_. *The OT problem can be interpreted as finding the joint distribution M that minimises the Kullback—Leibler divergence from* exp(*− C/ε*) *while satisfying the marginal conditions in* (6), *that is, matching with the observed distributions. This problem of matching an initial distribution with a target distribution under the assumption of some prior probabilities for the particle movements is known as the Schrödinger bridge problem [43, 19, 61]*.

### A.3 Model identification

The cost function can be defined for a known dynamical system, but the goal here is to use the OT approach in system identification, that is, determining the matrix *A* and the constant load vector *b*. We define the dynamical system as the solution to the minimisation problem

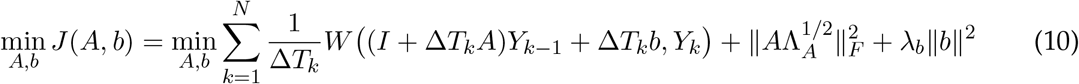

where the diagonal matrix Λ_*A*_ and *λ*_*b*_ are regularisation parameters and *W* is the entropy-regularised optimal transport cost defined in (6) with 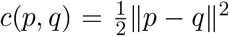 as the cost function. Say matrix 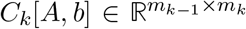contains the costs as defined in the previous section for measurements in *Y*_*k−*1_ and *Y*_*k*_. That is, the element (*i, j*) of this matrix is obtained by (9) where *x*_0_ is the *i*^th^ column of *Y*_*k−*1_ and *x*_1_ is the *j*^th^ column of *Y*_*k*_. Then expanding the definition of the optimal transport cost *W* results in a combined optimal transport and system identification problem:

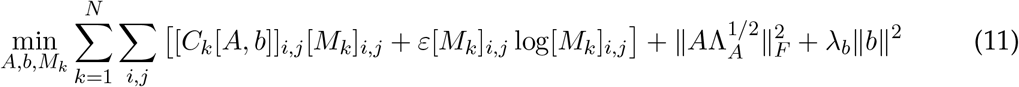

subject to *M*_*k*_𝟙 = 𝟙 and 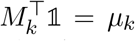. With this definition of the marginal distributions, each propagated cell is assumed to have mass one, whereas the masses of the cells in the target distribution are scaled. Usually the scaling is simply 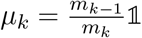, except in the cases with branching dynamics (see Section A.5). The combined cost function is strongly convex (quadratic) with respect to *A* and *b* when the transport plans *M*_*k*_ are fixed. Conversely, when *A, b* are fixed, the problem is convex (in fact strongly convex thanks to the entropy regularisation) with respect to the transport plans *M*_*k*_. Unfortunately, however, the componentwise convexity does not imply joint convexity. Indeed, the cost function in (11) may have several local minima. The componentwise convexity property nevertheless motivates a coordinate-descent type algorithm for solving the problem. Since the cost function is quadratic for the variables *A* and *b*, it can be solved analytically for fixed *M*_*k*_. When *A, b* are, in turn, fixed, (11) becomes a standard entropy-regularised OT problem and it can be solved efficiently with Sinkhorn iterations. The full problem is solved by alternating between these steps until convergence, as illustrated in Figure 1.

Note that with the transport plans *M*_*k*_ fixed, the regression problem (11) has 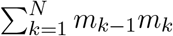 (degenerate) data points. As shown in Supplementary note 1, the problem can be reduced into a smaller and better explicable regression problem by defining a target point at time *T*_*k*_ for each cell *x*_*i*_(*k* − 1) measured at time *T*_*k−*1_ as a weighted average of the cells measured at time *T*_*k*_. The weights are given by the *i*^th^ row of the transport plan *M*_*k*_ (that sums up to one due to the marginal constraint *M*_*k*_𝟙 = 𝟙). With matrix notation, the target points for all cells in the matrix *Y*_*k−*1_ are given by 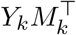. Moreover, these target points can be used to estimate derivatives for the cells (corresponding to RNA velocity [39]) by a difference quotient. Define a matrix containing all measured cells from time points *T*_0_ to *T*_*N−*1_ — augmented by a row of ones to include *b* in the same regression

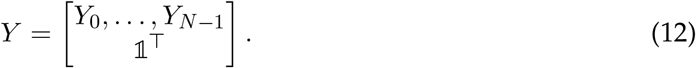

Their estimated derivatives are then given by

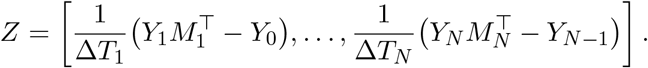

Then the augmented matrix [*A, b*] minimising (11) can be solved row-by-row from linear regression problems

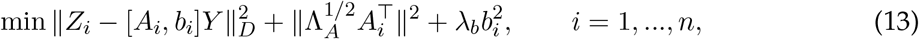

where *Z*_*i*_ and *A*_*i*_ are the *i*^th^ rows of matrices *Z* and *A*, and *D* is a diagonal matrix with diagonal elements 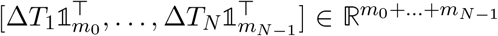. Note that the regression problem (13) has 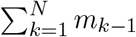data points instead of 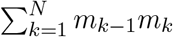 of the original problem (11).

To reduce the effect of noise in the data, for the solution of the OT problem, the dimension of the space is reduced by means of principal component (PC) reduction. The dimension of the PC space is 0.9*n* (rounded) but at most 100. Note, however, that this dimension reduction is applied only in the calculation of the costs in (9), but the identification of the dynamical system is done in the full *n*-dimensional space.

### A.4 Gene regulatory network inference

The variable selection step used for GRN inference is based on a greedy forward-backward sweep (inspired by [70] to which we also refer for details on the greedy approach) applied on the regression problem (13) one target gene at a time. As discussed in [70], the forward greedy approach may make mistakes that are not corrected and a backward greedy algorithm is more likely to work better. However, in high-dimensional problems, the backward algorithm may fail, if all possible regressors are allowed to be active at the same time. To initialise the backward greedy algorithm, the initial set of regressors is selected based on a combination of genegene correlations and a forward greedy algorithm. First, the regulators with high (positive or negative) correlation with the target gene are chosen in the regressor set (top *N*_reg_ *−* 5 genes), including the target gene itself, and the constant load term *b*_*i*_. To complement the set, a forward greedy algorithm is applied. At every step, the regressor yielding a highest decrease in the cost function is added to the regressor set. The forward sweep is continued until *N*_reg_ regressors are included. The backward greedy algorithm is then carried out, at every step removing the regressor that yields the smallest increase to the cost function. The exceptions are the gene itself and the constant load terms that are always kept in the regressor set. The backward greedy steps are continued until only these fixed regressors remain.

The method output is a confidence score for each link. During the backward phase, when there are *p* active regressors (not including the constant load terms), the cost function increments are collected in a vector Δ*J ∈* ℝ^*p*^ which is then normalised by dividing with max Δ*J* . Then, the link whose removal would yield the highest cost function increase, has value 1 in this vector. The final score of a particular link is the average of these scaled cost function increments calculated over those regressor sets where the link was active. Notice that with this scoring scheme, those links that at are never included in the active regressor set will get score zero. This scoring scheme has been developed with the aim to have values between zero and one for all genes, and to have values that are not too dispersed.

Note that for problems with small enough dimension, *n ≤ N*_reg_, only the backward greedy algorithm is applied starting from a full regressor set.

### A.5 Accounting for branching dynamics

Branching dynamics are a profoundly nonlinear phenomenon, and a linear model class is not ideal for dealing with it. A stable linear system (4) can only have one steady state, given by *x* = *− A*^*−*1^*b*. To include branching dynamics and enable multiple steady states, different branches are assigned to different constant vectors *b* in (4). Moreover, branching is taken into account in the OT problem by adjusting the weights of the cells *μ*_*k*_ and the costs *C*_*k*_ in (11). The goal is to discourage mass transport between branches in the solution of the OT problem.

The adjustment of the weights is done as follows. Say each measurement matrix *Y*_*k*_ is accompanied by a matrix 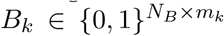 where *N*_*B*_ is the number of branches. This is an indicator matrix whose columns contain 0, 1 -valued variables indicating to which branch(es) the corresponding cell belongs to. Denote by *m*_*b*_(*k*) the sum of the *b*^th^ row of the matrix *B*_*k*_. Then, define *B*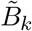 as the normalised indicator matrix, where each column of *B*_*k*_ is divided by the column sum. The idea is that cells belonging to multiple branches have their mass (in the sense of the OT problem) equally divided between branches. In case cells are only allowed to belong to one branch, it holds that 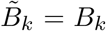. Denote by 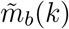 the sum of the *b*^th^ row of 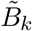 . Then the mass distribution *μ*_*k*_ of the cells of the *k*^th^ time point for the OT problem (6) is given by the column sum of diag 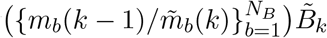 .

This scaling is done to enable the existence of a solution *M*_*k*_ that does not mix cells between branches. To encourage such solution, the cost matrix entries [*C*_*k*_[*A, b*]]_*i,j*_ are scaled by a factor higher than one when cells *i* and *j* do not belong to the same branch. This factor has a default value 2, but it can be changed by the user. To identify cell pairs between two time points that do not belong to the same branch, it suffices to find zeros of the matrix 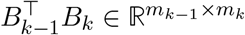.

Note that in this setup, one cell can belong to multiple branches (e.g., cells before the branching has taken place), although in the output of Slingshot, each cell is assigned to exactly one branch.

### A.6 Inference of perturbation targets

Perturbations are dealt with by introducing an additional vector *b* into the dynamics (4) in the variable selection phase of the method, but only to the dataset with the perturbation. That is, a line is added to the concatenated matrix *Y* defined in (12). This line has zeros in positions corresponding to the control experiment, and ones in positions corresponding to the perturbation experiment. This term is then considered as a possible regulator exactly as any gene, yielding an additional column into the method output matrix of confidence values of link existence.

### A.7 Inference of mutation effects

Mutations are handled by introducing a new line into the concatenated matrix *Y* defined in (12). This line has zeros in positions corresponding to the control experiment, and the expression values of the mutated gene in positions corresponding to the mutation experiment. Unlike with perturbations, this line is not used as an independent candidate regulator, but instead, it is considered an active regressor whenever the mutated gene is active. Effectively this means that the column of the *A*-matrix corresponding to the control and mutation datasets are allowed to be different. The introduction of a new row in the *Y* matrix also means that a new column is introduced in the *A*-matrix. This additional column is the difference of the *A*-matrix column corresponding to the mutation dataset compared to the control dataset. The backward greedy phase of the variable selection is then complemented with an additional step for evaluating mutation targets. If the mutated gene is an active regressor, the mutation is scored by calculating the cost function increment for removing only the additional mutation-row from the active regressor set (note that the mutated gene itself is scored by removing both the row corresponding to the mutated gene, and the additional row with zeros in places corresponding to the control experiment). This cost increment is then augmented into the Δ*J* vector and the method then proceeds as described in Section A.4. This results in an additional column in the method output matrix of confidence values of link existence.

### A.7 Lineage tracing and branch labeling

The optimal transport maps *M*_*k*_ obtained from (11) can be used to reconstruct cell trajectories, and to identify ancestor and descendant cells across measurement times. The columns (scaled to sum up to one) of the transport plan matrix *M*_*k*_ contain information about the propagated ancestors, while the rows (scaled) tell about the propagated descendants. It is possible to give a probabilistic interpretation to the scaled rows and columns of the transport matrix. That is, 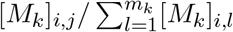 gives the probability that cell *i* measured at time *T*_*k−*1_ would be the ancestor of cell *j* measured at time *T*_*k*_, and 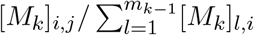 gives the probability that cell *j* would be the descendant of cell *i*. To identify descendant and ancestor cells across multiple time transitions, one can simply multiply the transport matrices corresponding to the transitions.

Among other things, the lineage tracing can be used to reconstruct branch labels that improve the performance of our method. This requires clustering the cells measured at the final time point where typically the clusters/branches can be identified relatively easily. We provide a function to then reconstruct branch labels for all time points using the transport matrices obtained from the method applied without using branch labels. The cluster labels should be given in the same format as the branch labels *B*_*k*_ described in Section A.5, that is, the label matrix is 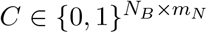 where *N*_*B*_ is the number of clusters/branches and *m*_*N*_ is the number of cells in the final time point. This matrix should contain exactly one non-zero entry on each column. The branch label for the last time point will be directly the cluster indicator given by the user, *B*_*N*_ = *C*. To reconstruct branch labels for a time point *k < N*, we calculate the transport map across multiple time points as a product 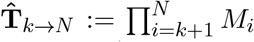 which is then normalised by the row sum to get 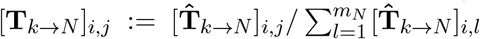. Each column of the matrix 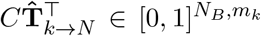 sums up to one. A column corresponds to a cell measured at time *T*_*k*_, and it gives the percentage of the descendants of the cell belonging to each branch (recall that the OT approach can split the cells such that each cell is associated with several past and future cells). The branch label matrix *B*_*k*_ is then constructed by inserting a 1 on each position where the corresponding entry of the matrix 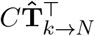 is greater than a threshold value, for which the default value is 1*/*(2*N*_*B*_). Note that with this approach, a cell can belong to multiple branches. If the user sets a threshold higher than 1*/N*_*B*_, it is possible that on some columns, no entry is higher than the threshold. In such case, a 1 is inserted in the column on the position with the highest entry to ensure that every cell is assigned to a branch. This approach is validated in Section 2.7 by comparing the branch assignments with those given by Slingshot.

### A.9 Additional features

Several experimental datasets can be combined into one inference problem. This, however, should only be done when the experiments are carried out on isogenic organisms.

While the link scoring described above always produces positive confidence values for link existence, the signs of the entries of the inferred *A*-matrix can be used to determine whether a regulatory link is an activation or an inhibition. GRIT can be requested to produce a signed list of confidence scores.

Parallelised computing is readily built-in in GRIT. The optimal transport problems are always solved independently for each time transition *T*_*k−*1_ *→ T*_*k*_, and therefore this step can be straightforwardly parallelised to up to *N* processors. The model update step, in turn, as well as the variable selection are done independently one target gene at a time, which can be done in parallel processes.

### A.10 Parameters

GRIT has few tuning parameters. As discussed above, the coefficient for entropic regularisation *ε* has a direct interpretation in terms of noise intensity in the data. However, noise intensity is not known, and, moreover, the real data are anyway not produced by linear discrete-time dynamics. The parameter is therefore selected to ensure robust solvability and stability of the Sinkhorn iterations used to solve the OT problem. Each time the OT problem is solved, as a preliminary step, the median of all entries in the cost matrix is calculated and *ε*_0_median(*C*) is used as the regularisation parameter. The coefficient *ε*_0_ has a default value 0.05, but it can be adjusted by the user.

The Tikhonov regularisation parameters Λ_*A*_ and *λ*_*b*_ in (10) are defined with the help of the matrix *Y* defined in (12) by calculating diag(*Y Y* ^*T*^), dividing by the total number of cells, and multiplying by the experiment time range *T*_*N*_ *− T*_0_ (or the sum of time ranges in case multiple experiments are concatenated). The resulting vector has dimension *n* + *N*_*B*_. The first *n* components multiplied by 0.01 are the diagonal entries of the matrix Λ_*A*_. The remaining *N*_*B*_ elements multiplied by 0.1 are used as the regularisation parameters for their respective *b* vectors corresponding to different branches. This regularisation is invariant to the time scale (results do not change if time unit is changed from days to hours, for example) and to the scales of different genes. Overall, the regularisation is rather mild, but it ensures stability of the solution even for low cell numbers and high dimension.

## B. Details on method application and result processing

### B.1 Datasets, preprocessing, and details of method application

To demonstrate that the method works as a tool for system identification in the framework in which it is designed, we formulate a 10-dimensional discrete-time linear system. This system is used to simulate single-cell data consisting of distributions at different time points. This system and the data generation process is detailed in Supplementary note 3. To demonstrate the effect of continuous-time dynamics, a continuous-time version is used to produce similar data. Further, a nonlinear version of the model is developed, where linear activations and inhibitions are replaced by nonlinear Michaelis–Menten kinetics, tuned in such way that the saturation limits have the same orders of magnitude as the gene expression levels in the linear case to ensure that nonlinearities do affect the system dynamics. Moreover, the noise processes in the nonlinear case are replaced by Langevin processes, which are more realistic models of noise in gene expression [23]. A simple model of protein dynamics is also included in the nonlinear model. However, the method only uses RNA concentration data, since protein concentrations are typically not measured together with RNA. To assure that the results obtained for the different systems are comparable, the systems are designed in such way that the linear system is a linearisation of the nonlinear system at the steady state point. Moreover, same noise process realisations are used for the systems (with the exception of the discrete time dynamics).

Method validation is done using the BEELINE benchmarking pipeline [55, 54]. The pipeline consists of three different types of datasets. The *synthetic* dataset consists of simulated data from six systems that are purpose-designed to reproduce certain qualitative behaviors, such as bifurcation, periodic dynamics, etc. For each system, ten datasets are simulated, and subsets with five different numbers of cells (100, 200, 500, 2000, or 5000) are provided for the inference tasks. This results in 300 tasks in total. The cases with 2000 and 5000 cells are used in the main comparison and the results for other cases are presented in supplementary material. The *curated* dataset consists of simulated data from four models from literature that have been created to mimic certain biological processes. Again, ten sets are simulated for each model, and for each of these replicates, three sets are created with different rates of dropouts in the data (dropout rates of 0%, 50%, and 70%). This results in 120 inference tasks in total. The case without dropouts is used in the main comparison and the results for other cases are presented in supplementary material. Finally, five real single cell RNA-Seq datasets are used for validation. One of the datasets corresponds to cell differentiation, and there are three different target cell types. The branching dynamics are identified with Slingshot, and the branches are treated as separate inference tasks. For each case, four different ways to select the genes of interest are used, resulting in 28 inference tasks in total.

GRIT requires data in batches of cells measured at different times, as is typical for a singlecell experiment. The simulated data of BEELINE do not follow this paradigm, but instead consists of cells whose simulation time has been randomly selected from some interval. We generated measured batches by ordering cells by their pseudotime (pseudotime by Slingshot is provided with the BEELINE data [54]), and then divided the data into eight or fifteen separate time points. The measurement time for each batch was the average of the pseudotimes of the cells in the batch. The real data in BEELINE consist of separate time points, except for three cases. In these three cases, we used the pseudotime (provided together with the data in [54]) to split the data into six time points.

Our biological objective is to look into genetic mechanisms of Parkinson’s disease (PD). In particular, we look into two datasets studying the effect of PD-associated mutations on the neuron development, namely a mutation of the *PINK1* gene [50] and a mutation on the *LRRK2* gene [68] (referred to as the *PINK1* and *LRRK2* datasets in the article). These datasets contain single cell RNA-seq data for the cellular differentiation process into dopaminergic neurons (DA). The *LRRK2* dataset monitored human patient specific induced pluripotent stem cell (iPSC)derived neuroepithelial stem cells’ (hNESCs) differentiation into DA. The mutant cells had the PD-associated *LRRK2*-G2019S mutation while the control cells were isogenic, corrected for the *LRRK2* mutation, and the samples were analysed on days 0, 10, 14, and 42 of the differentiation processes. The *PINK1* dataset monitored human patient specific induced pluripotent stem cells’ (iPSCs) differentiation into DA. The mutant cells had the PD-associated *PINK1*-I368N mutation while the control cells were from age-sex matched controls, and the samples were analysed on days 0, 6, 15, and 21 of the differentiation processes, and on an additional day 10 for the control cells.

For the *LRRK2* data, cells with less than 1000 reads were filtered out and only genes included in all data files (time points) were kept. In the *PINK1* dataset, the released data have fewer cells overall but the quality seems better, as all cells have more than 1000 reads, except for time points 10 and 21 in the control case. In fact, time point 10 has average read count 2804 and time point 21 has 1292 compared to the average 12717 for the remaining three time points. Due to this drastic discrepancy, these two time points are excluded from the analysis. Then, genes that were expressed in less than 10% of the cells were filtered out. The resulting *LRRK2* dataset had 6507 genes (down from 16662) with 3838 cells in total for the control case and 4403 cells in the mutant case. The filtered *PINK1* dataset had 8756 genes (down from 18097) with 2518 cells in the control case and 1977 cells in the mutation case. Wasserstein distances were calculated individually for all 1D distributions of different genes between consecutive time points (in 1D, the Wasserstein distance is the *L*^1^ norm between the cumulative distribution functions) and added up over all time points and both conditions. The genes were then sorted based on this dynamical variability, and 500, 1000, 1500, or 2000 most highly varying genes were given to GRIT. In the *LRRK2* dataset, *LRRK2* itself is only expressed in a handful of cells, and it is filtered out in the first phase. It can hardly be expected that any differences in regulations by *LRRK2* could be inferred based on a handful of reads. This experiment is therefore treated as a perturbation dataset rather than a mutation dataset, and it is treated as described in Section A.6. *PINK1* is better expressed, but it is not among the 2000 most highly varying genes. Therefore *PINK1* is manually added to the list of genes after the dimension reduction based on gene variability. GRIT was then run in two ways, regarding the second experiment either as a mutation experiment (treated as described in Section A.7), or as a perturbation experiment (treated as described in Section A.6). Both results are reported.

### B.2 Performance evaluation

The performance evaluation of GRIT uses standard scores, namely the Area Under the Receiver Operating Characteristics curve (AUROC), and the Area Under the Precision–Recall curve (AUPR). The AUPR score should be regarded as the primary metric due to class imbalance between existing and non-existing links (network sparsity) and because emphasis should be put on correctness of high-confidence predictions.

For a straightforward comparison, in the context of the BEELINE benchmarking pipeline, we use the performance metrics proposed in BEELINE. Since the AUPR values depend on the sparsity level of the network, the AUPR is used as a ratio of the AUPR of the inference results and the expected AUPR for random matrix. Then, AUPR-ratio 1 means a result comparable to random guessing. An additional metric is the Early Precision Ratio (EPR), which is also used in BEELINE. To calculate the EPR, *k* links with highest confidence assigned by a method are checked, and the share of true positive links is calculated. This share is then divided by the sparsity level of the network. This scaling means that an EPR of 1 corresponds to performance equivalent of random guessing. The value of *k* is chosen as the number of links in the ground truth network. In the main text, only summary results are shown on the BEELINE benchmark based on method ranking in different tasks. Full results are shown in the Supplementary material. The summary results are based on ranking the methods in each task. For synthetic dataset, AUPR-ratios for the six systems are used for this ranking (corresponding to [55, Fig. 2]). For the curated dataset, ranks were separately determined using the AUPR-ratio and the EPR (corresponding to [55, Fig. 4]). For the RNA-Seq data, EPRs for each seven tasks were used for ranking with the STRING and Non-specific ChIP-Seq networks used as ground truths in the cases TFs + 500 and 1000 genes (corresponding to [55, Fig. 5]). The results obtained using the Celltype-specific ChIP-Seq as the ground truth (and the lof/gof network for mESC) seemed to bear almost no correlation with the predictions of any of the evaluated methods, and these were therefore excluded from the summary results. The summary results for the RNA-Seq data are then based on 28 ranks (7 tasks, two different sizes, and two different ground truths).

### B.3 Pathway enrichment analysis for the *PINK1* and *LRRK2* results

As described above, for the analysis of the *PINK1* and *LRRK2* datasets, subsets of genes were first selected based on their dynamical variability, and then GRIT was run on these subsets. Of particular interest in the results are the genes that GRIT identifies as potential direct downstream effects of the mutation. GRIT outputs a scored list of all genes in the analysis where the score indicates the confidence for the gene being a perturbation target. To gain insight into these results, pathway enrichment analysis is then applied to this list.

To avoid bias due to the selection of genes included in the analysis and to avoid the need to select a threshold for the results, the following approach was taken. The pathways in the KEGG database [34] were used in the enrichment analysis. First, we determined the intersections of all terms in the KEGG database and the set of genes included in the analysis (using the g:profiler tool [37]). The genes included in the analysis were ordered by the confidence score for being a perturbation target according to GRIT. We determined the ranks of all genes for each KEGG term on the ordered list. Subsequently, we performed a Kolmogorov–Smirnov (KS) test with the null hypothesis that these ranks were uniformly distributed. One-tail KS test was used with the alternative hypothesis that the ranks originate from a cumulative distribution that is greater than the cumulative distribution of the uniform distribution to identify only pathways that are enriched in the high score genes. To correct for multiple hypothesis testing, p-values were adjusted by the Benjamini–Hochberg false discovery rate approach [10].

