## Supplementary material for "Gene regulatory network inference from single-cell data using optimal transport"

### Supplementary note 1:
### Reduction of the regression problem

The optimisation problem formulated in Equation (11) of the main text is a quadratic problem with respect to $A, b$ when the transport plans $M_k$ are fixed. To solve the quadratic problem, let us calculate the derivative of one entry in the cost matrix (using again $x_0$ and $x_1$ as generic vectors as in Equation (9) of the main text)

$$\frac{d}{dA}C_k[A,b] = \frac{d}{dA}\frac{1}{2\Delta T_k}\left\|x_1 - \big(I+\Delta T_k A\big)x_0 - \Delta T_k b\right\|^2 = -\big(x_1 - \big(I+\Delta T_k A\big)x_0 - \Delta T_k b\big)x_0^\top.$$

Then, denoting by $x_j(k)$ the cell $j$ measured at time $T_k$, it holds that

$$\begin{aligned}&\frac{d}{dA}\sum_{i=1}^{m_{k-1}}\sum_{j=1}^{m_k}[C_k[A,b]]_{i,j}[M_k]_{i,j}\\ &= -\sum_{i=1}^{m_{k-1}}\sum_{j=1}^{m_k}[M_k]_{i,j}\big(x_j(k) - \big(I+\Delta T_k A\big)x_i(k-1) - \Delta T_k b\big)x_i(k-1)^\top.\end{aligned}$$

Here anything multiplying $A$ does not depend on $j$, and therefore the marginal condition $M_k\mathbb{1} = \mathbb{1}$ gives

$$\begin{aligned}&\sum_{j=1}^{m_k}[M_k]_{i,j}\big[\big(I+\Delta T_k A\big)x_i(k-1) + \Delta T_k b\big]x_i(k-1)^\top\\ &= \big[\big(I+\Delta T_k A\big)x_i(k-1) + \Delta T_k b\big]x_i(k-1)^\top\end{aligned}$$

and therefore

$$\begin{aligned}&\frac{d}{dA}\sum_{i=1}^{m_{k-1}}\sum_{j=1}^{m_k}[C_k[A,b]]_{i,j}[M_k]_{i,j}\\ &= -\Delta T_k\sum_{i=1}^{m_{k-1}}\left[\frac{1}{\Delta T_k}\left(\sum_{j=1}^{m_k}[M_k]_{i,j}x_j(k) - x_i(k-1)\right) - Ax_i(k-1) - b\right]x_i(k-1)^\top.\end{aligned}$$

Similar calculation can be done for the derivative with respect to $b$.

Basically this means that when solving the optimal $A$ and $b$, this is equivalent to a regression problem where for each cell $x_i(k-1)$ measured at time $T_{k-1}$, a target point at time $T_k$ is defined as a weighted average of the cells measured at time $T_k$, where the weights are given by the $i^{\text{th}}$ row of the transport plan $M_k$. Moreover, this target point can be used to estimate a derivative for $x_i(k-1)$ by a difference quotient. That is, define a matrix containing all measured cells from time points $T_0$ to $T_{N-1}$ (augmented by a row of ones)

$$Y = \begin{bmatrix}Y_0,\ldots,Y_{N-1}\\ \mathbb{1}^\top\end{bmatrix}$$

and a matrix containing their estimated derivatives

$$Z = \left[\frac{1}{\Delta T_1}\big(Y_1M_1^\top - Y_0\big),\ldots,\frac{1}{\Delta T_N}\big(Y_NM_N^\top - Y_{N-1}\big)\right].$$

Then the augmented matrix $[A,b]$ can be solved row-by-row from linear regression problems $\min\|Z_i - [A_i,b_i]Y\|_D^2$ for $i = 1,...,n$, where $Z_i$ and $A_i$ are the $i^{\text{th}}$ rows of matrices $Z$ and $A$, and $D$ is a diagonal matrix with diagonal elements $[\Delta T_1\mathbb{1}_{m_0}^\top,\ldots,\Delta T_N\mathbb{1}_{m_{N-1}}^\top] \in \mathbb{R}^{m_0+\ldots+m_{N-1}}$

### Supplementary note 2:
### Consistency theorem for the method

In the absence of branching dynamics, and with $m$ cells measured at each time point, the method is based on minimisation of the cost function

$$\min_{A\in\mathbb{R}^{n\times n},b\in\mathbb{R}^n} \sum_{k=1}^{N} \frac{1}{2\Delta T_k} W\big((I+\Delta T_k A)Y_{k-1} + \Delta T_k b, Y_k\big) + \frac{1}{m}\|A\Lambda_A^{1/2}\|_F^2 + \frac{1}{m}\lambda_b\|b\|^2 \tag{1}$$

where $W(P,Q)$ is the discrete optimal entropy-regularised transport cost defined between point clouds $P=[p_1,...,p_{m_P}]$ and $Q=[q_1,...,q_{m_Q}]$ by

$$W(P,Q) = \min_{M\in\mathbb{R}^{m_P\times m_Q}} \sum_{i=1}^{m_P}\sum_{j=1}^{m_Q} \left[M_{i,j}\|p_i-q_j\|^2 + 2\varepsilon M_{i,j}\log(M_{i,j})\right] + 2\varepsilon\log(m_P m_Q) \tag{2}$$

subject to $M\mathbb{1} = \frac{1}{m_P}\mathbb{1}$ and $M^\top\mathbb{1} = \frac{1}{m_Q}\mathbb{1}$. Note that here the marginal constraints are slightly different from the main text. The different definition is compensated by scaling by $1/m$ in (1), which makes cost function (1) equivalent to the cost function (3) in the main text (up to multiplication by $1/m$ to prevent the cost from tending to infinity as $m$ increases) when there is no branching dynamics. Similarly, the term $2\varepsilon\log(m_P m_Q)$ in (2) was not a part of the definition in the main text, but it is constant with given data. These changes are required to make the discrete cost consistent with the continuous cost appearing in the proof below where the entropy regularisation involves a Kullback–Leibler divergence. Without this term, the entropy term tends to $-\infty$ when $m_P, m_Q \to \infty$. In this section we make the notational distinction between the cost function variables $(A,b)$ and the true system parameters $(\bar{A},\bar{b})$ that are assumed to have generated the observed data.

Assume the data is produced by a discrete-time system

$$x(k\Delta t) = (I+\Delta t\bar{A})x((k-1)\Delta t) + \Delta t\bar{b} + w_k \tag{3}$$

where $w_k \sim \mathcal{N}(0,\varepsilon I)$, $\varepsilon > 0$, and $w_k \perp w_j$ if $j\neq k$, and the initial state is a sample from a normal distribution $x(0)\sim\mathcal{N}(m_0,P_0)$. Assume that each measured cell is an independent realisation of this process. Each cell measured at time $k\Delta t$ is then an independent sample from a normal distribution $\mathcal{N}(m_k,P_k)$ where $m_k$ and $P_k$ satisfy the moment equations

$$\begin{cases} m_k = (I+\Delta t\bar{A})m_{k-1} + \Delta t\bar{b} \\ P_k = (I+\Delta t\bar{A})P_{k-1}(I+\Delta t\bar{A}^\top) + \varepsilon I. \end{cases} \tag{4}$$

Here it is assumed that each gene has the same input noise intensity. General noise covariance $\varepsilon R > 0$ can be handled by using $R^{-1}$ as a weight matrix in the Euclidean norm in (2).

**Theorem 1.** *Assume that the data consists of independent samples from a linear discrete-time system (3) and that at least three consecutive time points ($k=0,1,2,...$) have been measured with the following assumptions on the means and covariances of the measured distributions:*

*(i) $P_0 > 0$ and $I+\Delta t\bar{A}$ is invertible;*

*(ii) The algebraic multiplicity of any eigenvalue of $P_0^{-1/2}P_1P_0^{-1/2}$ does not exceed two;*

*(iii)* $P_0^{-1/2}(m_1 - m_0)$ *is not orthogonal to any eigenvector of* $P_0^{-1/2} P_1 P_0^{-1/2}$*. In case an eigenvalue has multiplicity two, then* $P_0^{-1/2}(m_1 - m_0)$ *is not orthogonal to the subspace spanned by the corresponding eigenvectors.*

*Assume further that the number of cells measured at each time point tends to infinity. Then the cost function (1) converges in* $L^1$ *(in stochastic sense) to a continuous cost function whose unique global minimiser is the true system* $(\bar{A}, \bar{b})$*, provided that the noise intensity* $\varepsilon$ *is used as the entropy regularisation parameter.*

Assumptions (i)–(iii) are needed for unique identifiability of the model, and they are precisely the assumptions in [2, Corollary 2.1]. If they are violated, there may exist several systems that can produce the observed data (in the case of normally distributed data). In this case, no method can be guaranteed to find the true system.

In the method, the regularisation constants $\Lambda_A$ and $\lambda_b$ do not scale up with the number of cells measured, whereas the transport cost does. Therefore their effect vanishes at the infinite-data limit, and the theorem holds for the method as such. However, the noise intensity $\varepsilon$ is never known in a real application and it is estimated from the data.

*Proof.* In the cost function (1), cells measured at time $k-1$ are propagated using a deterministic model, but they are compared to cells measured at time $k$ that have evolved according to stochastic dynamics (3). This discrepancy is accounted for by the entropy regularisation, which corresponds exactly to the noise in the dynamics of the true data.

The proof consists of three parts. In the first part, it is established that as the number of cells tends to infinity, the discrete entropy-regularised transport cost defined for point clouds sampled from probability distributions converges to the continuous transport cost defined for the distributions. In the second part, we show that for given normal distribution $\mathcal{N}(m_a, P_a)$ where $P_a > \varepsilon I$, the normal distribution $\mathcal{N}(m_b, P_b)$ with $m_b = m_a$ and $P_b = P_a - \varepsilon I$ is the unique minimiser of the entropy-regularised transport cost between $\mathcal{N}(m_a, P_a)$ and $\mathcal{N}(m_b, P_b)$. In the third part, relying on our earlier identifiability result, it is shown that the true system $(\bar{A}, \bar{b})$ is the unique system producing minimal entropy-regularised transport cost.

**Part 1.** Let $\alpha_N = \{a_1, ..., a_N\}$ and $\beta_N = \{b_1, ..., b_N\}$, be collections of independent samples from two sub-Gaussian probability distributions, $a_j \sim \alpha$ and $b_j \sim \beta$. In [5, Theorem 2], it is shown that the discrete entropy-regularised optimal transport cost $W(\alpha_N, \beta_N)$ defined in (2) converges with rate $N^{-1/2}$ in $L^1$ as $N \to \infty$ to the continuous transport cost (using the notation of [7] which we also refer to for details on the continuous problem)[1]

$$\mathcal{L}^\varepsilon(\alpha, \beta) = \min_{\pi \in \mathcal{U}(\alpha,\beta)} \int_{\mathcal{X}\times\mathcal{Y}} \|x - y\|^2 \pi(x,y) dx\, dy + 2\varepsilon \mathrm{KL}(\pi | \alpha \otimes \beta)$$

where $\mathcal{U}(\alpha, \beta)$ is the set of feasible transport plans, that is, probability distributions in $\mathbb{R}^n \times \mathbb{R}^n$ satisfying the marginal constraints $\int \pi(x,y)dy = \alpha(x)$ and $\int \pi(x,y)dx = \beta(y)$. The stochastic $L^1$ convergence means $\mathbb{E}\big(|W(\alpha_N, \beta_N) - \mathcal{L}^\varepsilon(\alpha, \beta)|\big) \to 0$ as $N \to \infty$.

**Part 2.** The proof of part 2 relies on a closed form expression of $\mathcal{L}^\varepsilon(\alpha, \beta)$ in case $\alpha$ and $\beta$ are normal distributions [4]:

$$\begin{aligned}\mathcal{L}^\varepsilon\big(\mathcal{N}(m_a, P_a), \mathcal{N}(m_b, P_b)\big) = \|m_a - m_b\|^2 + \mathrm{Tr}(P_a) + \mathrm{Tr}(P_b) - \mathrm{Tr}(D) \\ + n\varepsilon(1 - \log(2\varepsilon)) + \varepsilon \log(\det(D + \varepsilon I))\end{aligned}$$

[1]As we only discuss Gaussian distributions, we write the continuous OT problem in terms of density functions instead of general probability measures.

where $D = \big(4P_a^{1/2} P_b P_a^{1/2} + \varepsilon^2 I\big)^{1/2}$. The task is to find $(m_b, P_b)$ that minimise the transport cost for given $(m_a, P_a)$ where $P_a > \varepsilon I$. Clearly, the minimising $m_b$ is given by $m_b = m_a$. It is easier to find the minimum initially as a function of $D$ rather than $P_b$. To perform this change of variables, the covariance $P_b$ is expressed as a function of $D$:

$$P_b = \frac{1}{4} P_a^{-1/2}(D^2 - \varepsilon^2 I)P_a^{-1/2}.$$

The mapping between $D \geq \varepsilon I$ and $P_b \geq 0$ is bijective. Collecting the terms from the transport cost that depend on $D$ yield the minimisation problem

$$\min_{D \geq \varepsilon I} \frac{1}{4}\mathrm{Tr}\big(P_a^{-1/2} D^2 P_a^{-1/2}\big) - \mathrm{Tr}(D) + \varepsilon \log(\det(D + \varepsilon I)). \tag{5}$$

The target function is differentiable with respect to $D$ and the zeros of the derivative can be solved as follows (we refer to [6] for the derivative calculations):

$$\begin{aligned} &\frac{1}{2} P_a^{-1} D - I + \varepsilon (D + \varepsilon I)^{-1} = 0 \\ \Leftrightarrow\ \ &\frac{1}{2} P_a^{-1} D^2 + \frac{\varepsilon}{2} P_a^{-1} D - D = 0 \\ \Leftrightarrow\ \ &D = 2P_a - \varepsilon I \end{aligned}$$

where the last equivalence holds since $D$ is invertible. The cost function tends to infinity when $\mathrm{Tr}(P_b) \to \infty$, and therefore the minimum must be obtained either at the boundary of the feasible set $\{D \geq \varepsilon I\}$, or at $D = 2P_a - \varepsilon I$ which would yield the desired result $P_b = P_a - \varepsilon I$.

It is next shown that there can not be even a local minimum on the boundary of the feasible set. A matrix on the boundary has the singular value decomposition $D = USU^\top$ where $U$ is orthogonal, $S$ is diagonal with $S_{j,j} \geq \varepsilon$, and $S_{i,i} = \varepsilon$ for some $i$. It will be shown that the derivative of the cost function with respect to $S_{i,i}$ is negative, meaning that there exists a matrix inside the feasible set that attains a lower cost value than the matrix on the boundary. It holds that $\mathrm{Tr}(D) = \sum_{j=1}^n S_{j,j}$ and so $\frac{d}{dS_{i,i}}\big(-\mathrm{Tr}(D)\big) = -1$. The last term in (5) can be written in terms of the decomposition by $\varepsilon \log(\det(D + \varepsilon I)) = \varepsilon \sum_{j=1}^n \log(S_{j,j} + \varepsilon)$ and so $\frac{d}{dS_{i,i}}\big(\varepsilon \log(\det(D+\varepsilon I))\big)\big|_{S_{i,i}=\varepsilon} = 1/2$. Finally, for the first term in (5), substituting $D^2 = US^2U^\top$, it holds that

$$\frac{d}{dS_{i,i}} \mathrm{Tr}\left(\frac{1}{4} P_a^{-1/2} U S^2 U^\top P_a^{-1/2}\right)\Bigg|_{S_{i,i}=\varepsilon} = \frac{\varepsilon}{2} U_i^\top P_a^{-1} U_i < \frac{1}{2}$$

where the inequality follows from $P_a > \varepsilon I$. Here $U_i$ denotes the $i^{\text{th}}$ column of $U$. It follows that at any point on the boundary of the feasible set, the transport cost decreases when moving to the interior of the feasible set in such way that $D$ moves to the direction $U_i U_i^\top$. Therefore $P_b = P_a - \varepsilon I$ must be the unique global minimum.

**Part 3.** Time point $k$ consists of (infinite number of) samples from the normal distribution $\mathcal{N}(m_k, P_k)$ where $m_k$ and $P_k$ satisfy the moment equations (4). Based on part 2 of the proof, the normal distribution that minimises $\mathcal{L}^\varepsilon\big(\mathcal{N}(m_k, P_k), \mathcal{N}(m, P)\big)$ is given by $m = m_k = (I + \Delta t \bar{A}) m_{k-1} + \Delta t \bar{b}$ and $P = P_k - \varepsilon I = (I + \Delta t \bar{A}) P_{k-1} (I + \Delta t \bar{A}^\top)$. Note that by assumption (i), $P_k > \varepsilon I$ for $k \geq 1$.

In the method, samples measured at time point $k-1$ stored in matrix $Y_{k-1}$ are propagated by the candidate model $(A, b)$ by $(I + \Delta t A) Y_{k-1} + \Delta t b$. Since these are samples from a normal distribution $\mathcal{N}(m_{k-1}, P_{k-1})$, then the propagated points are samples from $\mathcal{N}\big((I + \Delta t A) m_{k-1} +$

$\Delta t b, (I + \Delta t A) P_{k-1} (I + \Delta t A^\top)\big)$. If $(A, b) = (\bar{A}, \bar{b})$, this is exactly the distribution minimising the transport cost to the distribution of measurements of the next time point. Based on [2, Corollary 2.1], the true system is the only possible model to produce the minimising distribution.

□

Part 2 of the proof shows that the entropy regularisation deconvolutes the noise $\varepsilon I$ out of the observations, whose covariance is $(I + \Delta t \bar{A}) P_{k-1} (I + \Delta t \bar{A}^\top) + \varepsilon I$. In the case of normal distributions, deconvolution is a simple subtraction of the noise covariance. This idea of entropy regularisation as noise deconvolution holds more generally and it has been explored in [9].

The theorem is a consistency result stating that the true system matrices are the unique global minimiser of the cost function (1). However, there is no guarantee that the method will necessarily converge to the global minimum instead of a local minimum. The method is initialised from $(A, b) = (0, 0)$, whereby in the first iteration of the algorithm the optimal transport problem is solved directly between the measured time points without any propagation through a candidate model. If the transport mapping obtained in this initial step is poorly representing true cell propagation, then there is a risk that the method does not find the global minimum. The further apart the time points are from each other and the more of the developmental dynamics are not observed, the higher is the risk of this happening.

### Supplementary note 3:
### Data generation for the experiments of Sections 2.2–2.5

#### Linear dynamics

Data is simulated either using discrete-time dynamics

$$x(k\Delta T) = (I + \Delta T A)x((k-1)\Delta T) + \Delta T b + w_k$$

where $w_k \sim \mathcal{N}(0, \Delta T Q)$, and measurement times are exactly $k\Delta T$ with $k = 0, 1, ..., N$, or using continuous-time dynamics

$$\frac{d}{dt}x(t) = Ax(t) + b + w(t)$$

where $w$ is a white noise process with covariance $Q$. The continuous-time system is simulated with Euler–Maruyama method using time step 0.01, which is well below the time scales of the measurements. To generate single-cell data, each cell is independently drawn from an initial distribution, and then propagated through the stochastic model for the time specified by the measurement time, at which the value of the trajectory is stored.

The system $A, b$ is given by

$$A = \begin{bmatrix} -1.5 & 0 & 0 & 0 & 0 & 0 & 0.4053 & 0 & 0 & 0.4053 \\ 0.2405 & -1.5 & -0.3269 & 0 & 0 & 0 & 0 & 0 & 0 & 0 \\ 0 & 0.4887 & -0.4 & 0 & 0 & 0 & 0 & 0 & 0 & 0 \\ 0 & 0 & 0.1509 & -1.5 & 0 & 0 & 0 & 0 & 0 & 0 \\ 0 & 0 & 0 & 0.6827 & -1.5 & 0 & 0 & 0 & 0 & 0 \\ 0 & 0 & 0.2658 & 0 & 0.2658 & -1.5 & 0 & 0 & 0 & 0 \\ 0 & 0 & 0 & 0 & 0 & 0.5555 & -2 & 0 & -0.4315 & 0 \\ 0 & 0 & 0 & 0 & 0 & 0 & 0.3080 & -0.8 & 0 & 0 \\ 0 & 0 & 0 & 0 & 0 & 0 & 0 & 0.7881 & -0.5 & 0 \\ 0 & 0 & 0 & 0 & 0 & 0 & 0 & 0 & 0.4675 & -1.2 \end{bmatrix}$$

$$b = \begin{bmatrix} 3.4750, & 2.4679, & 1.1554, & 2.7959, & 1.0879, & 2.4127, & 1.3674, & 0.4087, & 3.5551 \end{bmatrix}^\top.$$

This system is based on the system used in [1]. However, to study the effect of nonlinearities in the system dynamics, a nonlinear system was first generated, which is described below. The linear system has been obtained by linearising the nonlinear system at the system's steady state. This way, the behaviours of the linear and nonlinear systems are highly similar. Time step $\Delta T = 0.4$ is used in the experiment and six time points are generated with 1000, 2000, 3000, 5000, 7500, and 10000 cells in each time point in the experiment in Section 2.2 and 600 cells in each time point in the experiments of Sections 2.3–2.5. The initial states of each cell are drawn from a normal distribution $\mathcal{N}(m_0, P_0)$ where each entry of $m_0$ is drawn from a uniform distribution $U(0, 2x_{\text{ss},j})$. The initial state covariance is randomised by the following procedure:

$$\begin{aligned} R_0 &= \text{randn}(10, 10) && \text{(randomisation)} \\ R_1 &= R_0 R_0^\top && (\rightarrow \text{symmetric, pos. def.}) \\ R_2 &= \text{diag}(R_1)^{-1/2} R_1 \text{diag}(R_1)^{-1/2} && \text{(normalisation)} \\ P_0 &= \text{diag}(m_0)^{1/2} R_2 \text{diag}(m_0)^{1/2} + 0.3I && \text{(scaling)} \end{aligned}$$

where randn$(m, n)$ produces an $m \times n$ matrix with entries independently drawn from $\mathcal{N}(0, 1)$.

### Nonlinear dynamics

To demonstrate the effect of nonlinearity in the system dynamics, we created a nonlinear system by replacing all activations in the network with saturating Michaelis–Menten kinetics, where the saturation levels were chosen in a way that nonlinearity has an effect on the dynamics. Moreover, the two negative elements in the $A$-matrix above were replaced by proper inhibitions in the model.

The dynamics of each gene are governed by the Langevin equation ([3])

$$\frac{dx_j}{dt} = -a_j x_j + f_j(x) + \kappa\big(\sqrt{a_j x_j}\, w_j + \sqrt{f_j(x)}\, v_j\big) \tag{6}$$

where $w_j$ and $v_j$ are independent white noise processes corresponding to uncertainty in RNA degradation and expression processes, respectively. The noise scaling coefficient $\kappa$ is related to the molecule numbers within a cell. The higher the molecule numbers are, the smaller is the effect of noise. If the state vector directly corresponds to actual molecule numbers, then $\kappa = 1$ should be used. We, however, use it as a tuning parameter. The functions $f_j$ are gene expression rates and are therefore always non-negative. Without the noise, any simulation that is initiated from non-negative values in $x(0)$ will remain non-negative. Since the Langevin-approximation of the chemical master equation may violate the non-negativity assumption, after every time step in the numerical simulation, possible negative values are set to zero. The functions $f_j$ for $j = 1, ..., 10$ are as follows:

$$f_j(x) = \frac{v_j + k_j x_{j-1}}{1 + c_j x_{j-1}}, \qquad \text{for } j = 3, 4, 5, 8, 9, 10,$$

$$f_1(x) = \frac{v_1 + k_1(x_7 + x_{10})}{1 + c_1(x_7 + x_{10})}, \qquad f_2(x) = \frac{v_2 + k_2 x_1}{(1 + c_2 x_1)(1 + d_2 x_3)},$$

$$f_6(x) = \frac{v_6 + k_6(x_3 + x_5)}{1 + c_6(x_3 + x_5)}, \qquad f_7(x) = \frac{v_7 + k_7 x_6}{(1 + c_7 x_6)(1 + d_7 x_9)}.$$

This nonlinear system has precisely the same regulatory interactions as the linear system above. That is, six genes (3,4,5,8,9,10) are activated by one gene ($x_{j-1} \to x_j$). Genes 1 and 6 are activated by two genes ($x_7, x_{10} \to x_1$ and $x_3, x_5 \to x_6$). Genes 2 and 7 have one activation and one inhibition ($x_3 \dashv x_2$, $x_1 \to x_2$ and $x_9 \dashv x_7$, $x_6 \to x_7$). The parameter values of the system are shown in the adjacent table.

To ensure that the linear and nonlinear dynamics would be as close to each other as possible (that is, nonlinearity being the only difference), same noise process realisations were used in the simulations. Since the nonlinear system (6) has two noise processes $w_j$ and $v_j$ per gene, for

**Table:** Parameters of the nonlinear model.

| Gene ($j$) | $a_j$ | $k_j$ | $v_j$ | $c_j$ | $d_j$ |
|---|---|---|---|---|---|
| 1 | 1.5 | 2.0991 | 0.4524 | 0.2015 | |
| 2 | 1.5 | 3.6469 | 0.5124 | 0.2318 | 0.6667 |
| 3 | 0.4 | 2.5882 | 0.4377 | 0.8641 | |
| 4 | 1.5 | 3.9127 | 0.0479 | 0.9068 | |
| 5 | 1.5 | 1.5617 | 0.3310 | 0.2069 | |
| 6 | 1.6 | 3.6369 | 0.1265 | 0.4267 | |
| 7 | 2.0 | 2.2910 | 0.3932 | 0.0411 | 0.4545 |
| 8 | 0.8 | 3.7228 | 0.4238 | 1.5239 | |
| 9 | 0.5 | 1.0370 | 0.1570 | 0.0628 | |
| 10 | 1.2 | 3.2482 | 0.3040 | 0.3638 | |

the linear dynamics, $K(w_j + v_j)$ was used as the noise process, where $K$ is a diagonal matrix whose $j^{\text{th}}$ diagonal entry is $\sqrt{a_j x_{\text{ss},j}}$ with $x_{\text{ss},j}$ being the steady-state value of gene $j$. Then the noise intensities of the linear and nonlinear systems coincide at the steady state (note that $a_j x_{\text{ss},j} = f_j(x_{\text{ss}})$).

### Protein dynamics

Typically the RNA-molecules do not act as transcription factors directly, but are first translated into proteins. The protein levels, however, are typically not measured, and computational methods directly fit models between RNA concentrations. To test the effect of this omission, we modified the nonlinear system by introducing a variable mimicking protein concentrations whose dynamics are governed by

$$\frac{d}{dt}p(t) = L(x(t) - p(t)), \qquad p(0) = x(0)$$

where $L$ is a diagonal matrix with strictly positive diagonal entries, that are drawn from $U(0.5, 1)$ separately for each replicate. In the RNA dynamics (6), $p$ replaces $x$ as the variable of the functions $f_j$. No noise is considered in the protein dynamics, since we only wished to study the effect of the delay introduced by protein dynamics while ignoring the effect of additional noise. The initial state $p(0) = x(0)$ is chosen following an assumption that the population is in a steady state before a perturbation is introduced, and the protein levels have converged to correspond to this steady state (note that if $x(t) = x_c$ is constant, then $p(t) \to x_c$ as $t \to \infty$).

With the protein concentrations included, the system's dimension is 20, but the method is applied on the 10-dimensional data of RNA concentrations, and the protein concentrations are ignored.

### Mutation and perturbation model

The method can be used to find targets of external perturbations (for example, a drug) or pathways affected by a mutation. To generate data for validating the approach, the nonlinear model was slightly modified. In the original model, no gene regulates more than two other genes. To have a more meaningful validation experiment for mutation effects, gene 9 (which already activates gene 10 and inhibits gene 7) is given a third regulation, namely an activation of gene 4. In addition, the three regulatory effects of gene 9 are modulated by coefficients $s_i \in [0, 1]$, for $i = 1, 2, 3$, simulating a partial loss of function. That is, functions $f_4$, $f_7$, and $f_{10}$ in the model are replaced by

$$f_4(x) = \frac{v_4 + k_4(x_3 + s_1 x_9)}{1 + c_4(x_3 + s_1 x_9)}, \quad f_7(x) = \frac{v_7 + k_7 x_6}{(1 + c_7 x_6)(1 + d_7 s_2 x_9)}, \quad f_{10}(x) = \frac{v_{10} + k_{10} s_3 x_9}{1 + c_{10} s_3 x_9}.$$

This model was then run always first with $s_1 = s_2 = s_3 = 1$ to simulate a system without any mutation. Then, another experiment was simulated with setting one or two of the three coefficients to either 0.5 or 0.75. The initial distribution for the control and mutation datasets were the same. Ten replicates for each gene combination were simulated, that is, altogether 30 replicates for each of the four cases (50% loss of function for one regulation, 50% loss for two regulations, 25% loss for one regulation, and 25% loss for two regulations).

To simulate the effect of perturbations, the basal transcription rates $v_j$ were modulated by coefficients $r_j \geq 1$. As with the mutation dataset, one experiment was simulated without the perturbation ($r_j = 1$ for all $j$), and then another experiment with a perturbation. Different number (one, two, or three) of genes were randomly chosen as perturbation targets, and the coefficients $r_j$ were increased to either 1.3 or 1.6 for the affected genes, corresponding to 30%

or 60% increase in the basal transcription rate. For each of the six combinations of number of affected genes and perturbation strengths, 20 replicates were simulated with different perturbation targets.

**Supplementary table 1:** Complete EPR results on the (log-transformed) BEELINE RNA-Seq data. The columns "TFs + 500 genes" and "TFs + 1000 genes" are comparable to [8, Fig. 5] and columns "500 genes" and "1000 genes" are comparable to [8, Suppl. fig. 8].

| Ground truth | Data | TFs+ 500 genes | TFs+ 1000 genes | 500 genes | 1000 genes |
|---|---|---|---|---|---|
| STRING | mHSC-E | 4.574 | 4.537 | 3.262 | 3.635 |
| | mHSC-L | 4.738 | 5.000 | 4.925 | 5.000 |
| | mHSC-GM | 6.119 | 6.409 | 4.912 | 5.790 |
| | mESC | 2.816 | 2.732 | 2.301 | 2.535 |
| | mDC | 1.459 | 1.629 | 1.283 | 1.160 |
| Nonspecific ChIP-Seq | mHSC-E | 3.159 | 3.039 | 1.679 | 2.429 |
| | mHSC-L | 2.054 | 2.182 | 1.831 | 2.182 |
| | mHSC-GM | 4.002 | 3.307 | 3.259 | 3.699 |
| | mESC | 2.443 | 2.936 | 1.730 | 2.076 |
| | mDC | 2.204 | 2.323 | 0.956 | 1.742 |
| Celltype-specific ChIP-Seq | mHSC-E | 1.005 | 1.002 | 1.009 | 1.001 |
| | mHSC-L | 1.023 | 1.010 | 1.025 | 1.010 |
| | mHSC-GM | 1.001 | 1.008 | 0.983 | 0.987 |
| | mESC | 1.033 | 1.012 | 1.031 | 1.043 |
| | mDC | 0.836 | 0.877 | 0.938 | 1.043 |
| lof/gof | mESC | 1.263 | 1.171 | 1.246 | 1.185 |
| STRING | hESC | 2.973 | 3.133 | 2.868 | 2.850 |
| | hHep | 2.048 | 1.921 | 1.763 | 2.062 |
| Nonspecific ChIP-Seq | hESC | 1.600 | 1.390 | 1.380 | 0.505 |
| | hHep | 1.337 | 1.240 | 1.826 | 1.827 |
| Celltype-specific ChIP-Seq | hESC | 1.082 | 1.125 | 1.070 | 0.937 |
| | hHep | 1.010 | 1.007 | 0.999 | 1.017 |

**Supplementary table 2:** Results of pathway enrichment analysis for the *LRRK2* dataset. The table shows statistically significant results (rank 1–13 where adjusted p-value $< 0.05$) as well as other terms with p-value $< 0.05$, KS $> 0.4$, and #genes $> 1$ to highlight some terms whose statistical significance is thwarted by the small term size. Here KS is the Kolmogorov–Smirnov statistic, which is then multiplied by $(\#genes)^{1/2}$ to get the adjusted KS statistic. Adjusted p-values are obtained by multiplying the p-value by the number of pathways tested (314), and then dividing by the rank of each pathway. The column "top200" indicates the percentage of genes of the term that are on top-200 on the list given by GRIT.

| Rank | KEGG term name | p_value | p_adj | KS_adj | KS | #genes | top200 |
|---|---|---|---|---|---|---|---|
| **1** | Oxidative phosphorylation | 1.347e-06 | 0.0002272 | 2.551 | 0.3266 | 61 | 21.3 |
| **2** | Ribosome | 1.447e-06 | 0.0002272 | 2.554 | 0.2855 | 80 | 31.3 |
| **3** | Thermogenesis | 2.224e-06 | 0.0002328 | 2.51 | 0.2958 | 72 | 19.4 |
| **4** | Coronavirus disease - COVID-19 | 3.503e-06 | 0.000275 | 2.471 | 0.268 | 85 | 31.8 |
| **5** | Diabetic cardiomyopathy | 3.063e-05 | 0.001924 | 2.244 | 0.2702 | 69 | 17.4 |
| **6** | Huntington disease | 5.556e-05 | 0.002907 | 2.189 | 0.2106 | 108 | 13.0 |
| **7** | Non-alcoholic fatty liver disease | 0.0001223 | 0.005485 | 2.082 | 0.2945 | 50 | 14.0 |
| **8** | Chemical carcinogenesis - reactive oxygen species | 0.0001719 | 0.006747 | 2.052 | 0.237 | 75 | 20.0 |
| **9** | Amyotrophic lateral sclerosis | 0.0005326 | 0.01858 | 1.921 | 0.1783 | 116 | 13.8 |
| **10** | Prion disease | 0.0006707 | 0.02106 | 1.89 | 0.1853 | 104 | 12.5 |
| **11** | Parkinson disease | 0.0008161 | 0.0233 | 1.865 | 0.177 | 111 | 12.6 |
| **12** | Retrograde endocannabinoid signaling | 0.001331 | 0.03483 | 1.785 | 0.2632 | 46 | 19.6 |
| **13** | Cell cycle | 0.001581 | 0.03819 | 1.76 | 0.2684 | 43 | 2.3 |
| **...** | | | | | | | |
| **17** | DNA replication | 0.02292 | 0.4234 | 1.309 | 0.414 | 10 | 0 |
| **19** | Base excision repair | 0.02697 | 0.4455 | 1.259 | 0.514 | 6 | 16.7 |
| **20** | Homologous recombination | 0.02838 | 0.4455 | 1.228 | 0.614 | 4 | 0 |
| **22** | Nucleotide excision repair | 0.03311 | 0.4477 | 1.23 | 0.465 | 7 | 0 |
| **23** | IL-17 signaling pathway | 0.03326 | 0.4477 | 1.235 | 0.4365 | 8 | 50.0 |
| **26** | Tyrosine metabolism | 0.04162 | 0.4863 | 1.126 | 0.796 | 2 | 50.0 |
| **27** | Pyruvate metabolism | 0.04223 | 0.4863 | 1.191 | 0.421 | 8 | 0 |

**Supplementary table 3:** Results of pathway enrichment analysis for the *PINK1* dataset. The table shows statistically significant results (rank 1–27 where adjusted p-value $< 0.05$) as well as other terms with p-value $< 0.05$, KS $> 0.4$, and #genes $> 1$ to highlight some terms whose statistical significance is thwarted by the small term size. Here KS is the Kolmogorov–Smirnov statistic, which is then multiplied by $(\#genes)^{1/2}$ to get the adjusted KS statistic. Adjusted p-values are obtained by multiplying the p-value by the number of pathways tested (316), and then dividing by the rank of each pathway. The column "top200" indicates the percentage of genes of the term that are on top-200 on the list given by GRIT.

| Rank | KEGG term name | p_value | p_adj | KS_adj | KS | #genes | top200 |
|---|---|---|---|---|---|---|---|
| **1** | Serotonergic synapse | 5.028e-05 | 0.01589 | 2.055 | 0.685 | 9 | 44.4 |
| **2** | Lipid and atherosclerosis | 0.0001547 | 0.02445 | 2.018 | 0.4512 | 20 | 30 |
| **3** | Oxytocin signaling pathway | 0.0003221 | 0.03393 | 1.937 | 0.4226 | 21 | 38.1 |
| **4** | Dopaminergic synapse | 0.0005839 | 0.03401 | 1.863 | 0.4166 | 20 | 40 |
| **5** | Estrogen signaling pathway | 0.0007266 | 0.03401 | 1.821 | 0.4703 | 15 | 40 |
| **6** | Long-term potentiation | 0.0008513 | 0.03401 | 1.765 | 0.5884 | 9 | 44.4 |
| **7** | Parathyroid hormone synthesis, secretion and action | 0.0009532 | 0.03401 | 1.741 | 0.6156 | 8 | 37.5 |
| **8** | Cholinergic synapse | 0.001019 | 0.03401 | 1.744 | 0.5814 | 9 | 44.4 |
| **9** | Circadian entrainment | 0.001019 | 0.03401 | 1.744 | 0.5814 | 9 | 66.7 |
| **10** | GnRH signaling pathway | 0.001146 | 0.03401 | 1.72 | 0.6081 | 8 | 50 |
| **11** | Adrenergic signaling in cardiomyocytes | 0.001381 | 0.03401 | 1.757 | 0.3833 | 21 | 38.1 |
| **12** | Salmonella infection | 0.001436 | 0.03401 | 1.778 | 0.2442 | 53 | 18.9 |
| **13** | Cocaine addiction | 0.001488 | 0.03401 | 1.607 | 0.8036 | 4 | 25 |
| **14** | Glucagon signaling pathway | 0.001507 | 0.03401 | 1.733 | 0.4333 | 16 | 25 |
| **15** | Alcoholism | 0.001739 | 0.03663 | 1.714 | 0.4286 | 16 | 37.5 |
| **16** | cGMP-PKG signaling pathway | 0.002064 | 0.04076 | 1.705 | 0.3634 | 22 | 27.3 |
| **17** | Vascular smooth muscle contraction | 0.002239 | 0.04161 | 1.678 | 0.4332 | 15 | 46.7 |
| **18** | Human immunodeficiency virus 1 infection | 0.002533 | 0.04448 | 1.682 | 0.3299 | 26 | 23.1 |
| **19** | Cortisol synthesis and secretion | 0.002697 | 0.04485 | 1.544 | 0.7721 | 4 | 50 |
| **20** | Inflammatory mediator regulation of TRP channels | 0.002869 | 0.04533 | 1.607 | 0.568 | 8 | 62.5 |
| **21** | Aldosterone synthesis and secretion | 0.003086 | 0.04563 | 1.606 | 0.5352 | 9 | 55.6 |
| **22** | Sphingolipid signaling pathway | 0.003237 | 0.04563 | 1.616 | 0.4666 | 12 | 8.3 |
| **23** | Pathways in cancer | 0.003321 | 0.04563 | 1.664 | 0.2048 | 66 | 19.7 |
| **24** | Amphetamine addiction | 0.003496 | 0.04603 | 1.596 | 0.5046 | 10 | 50 |
| **25** | Viral carcinogenesis | 0.003763 | 0.0464 | 1.633 | 0.2761 | 35 | 14.3 |
| **26** | Pathogenic Escherichia coli infection | 0.003817 | 0.0464 | 1.634 | 0.2616 | 39 | 10.3 |
| **27** | Olfactory transduction | 0.004108 | 0.04808 | 1.536 | 0.6272 | 6 | 66.7 |
| **...** | | | | | | | |
| **31** | Salivary secretion | 0.01045 | 0.1039 | 1.418 | 0.5359 | 7 | 57.1 |
| **32** | Glutamatergic synapse | 0.01076 | 0.1039 | 1.414 | 0.5343 | 7 | 57.1 |
| **34** | Antigen processing and presentation | 0.01118 | 0.1039 | 1.427 | 0.4512 | 10 | 30 |
| **35** | Growth hormone synthesis, secretion and action | 0.01199 | 0.1083 | 1.411 | 0.4703 | 9 | 22.2 |
| **36** | Long-term depression | 0.01293 | 0.1135 | 1.377 | 0.562 | 6 | 33.3 |
| **37** | Morphine addiction | 0.01428 | 0.1219 | 1.35 | 0.6036 | 5 | 60 |
| **41** | Tuberculosis | 0.01821 | 0.1404 | 1.338 | 0.4731 | 8 | 25 |
| **42** | Phosphatidylinositol signaling system | 0.01884 | 0.1417 | 1.318 | 0.5382 | 6 | 33.3 |
| **45** | Renin secretion | 0.02119 | 0.1488 | 1.299 | 0.5305 | 6 | 50 |
| **49** | Pantothenate and CoA biosynthesis | 0.02654 | 0.1712 | 1.184 | 0.8371 | 2 | 50 |
| **55** | IL-17 signaling pathway | 0.0349 | 0.2005 | 1.194 | 0.5971 | 4 | 50 |
| **58** | C-type lectin receptor signaling pathway | 0.03736 | 0.2035 | 1.209 | 0.4568 | 7 | 14.3 |
| **63** | Insulin secretion | 0.04738 | 0.2376 | 1.164 | 0.44 | 7 | 42.9 |

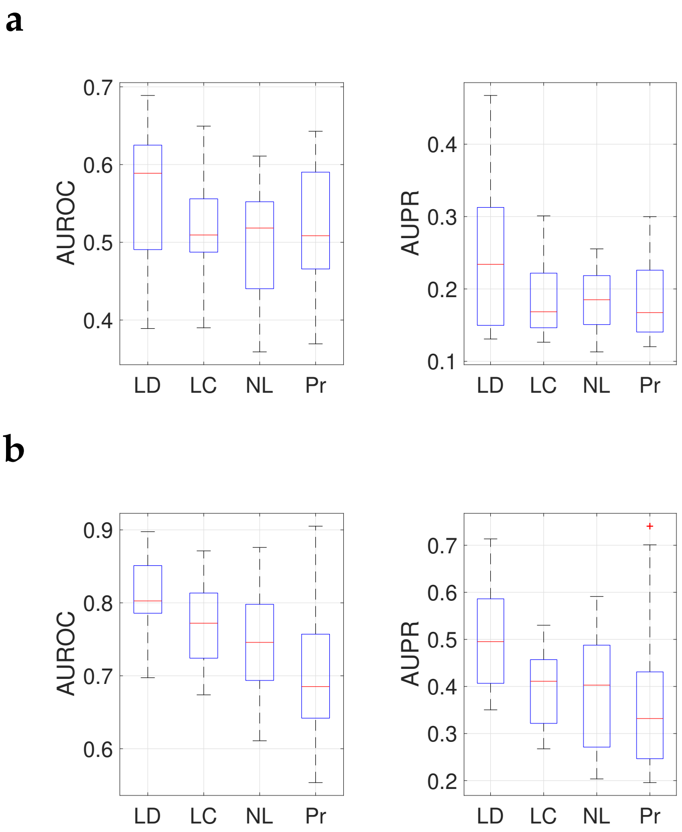


**Supplementary figure 1:** **a:** Results on the experiment of Section 2.3 by applying GRIT on bulk data, where each cell's expression vector has been replaced by the mean of the corresponding time point. **b:** Results on the experiment of Section 2.3 calculated from the absolute values of the $A$-matrix entries. These plots are directly comparable with Figure 2b in the main text.

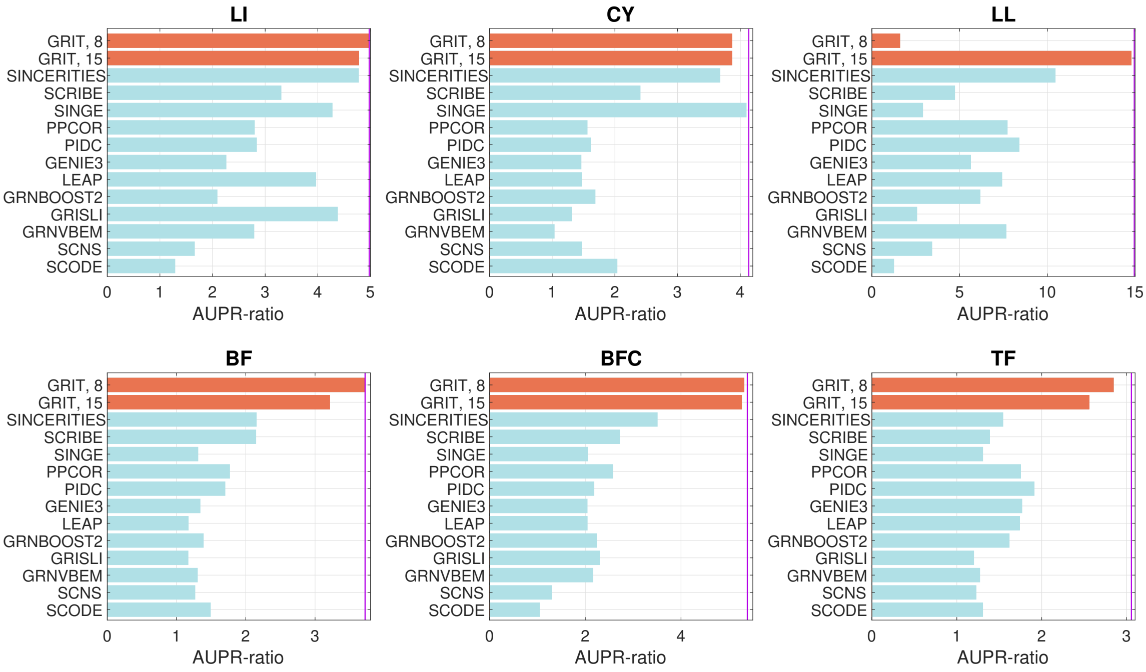


**Supplementary figure 2:** Detailed results of the BEELINE synthetic dataset. Bars show the median AUPR-ratios for cases with 2000 or 5000 cells (corresponding to [8, Fig. 2]). Results for the other methods are taken from the source file for [8, Fig. 2]. The purple line indicates the score for perfect reconstruction (the inverse of **E**(AUPR) for random networks).

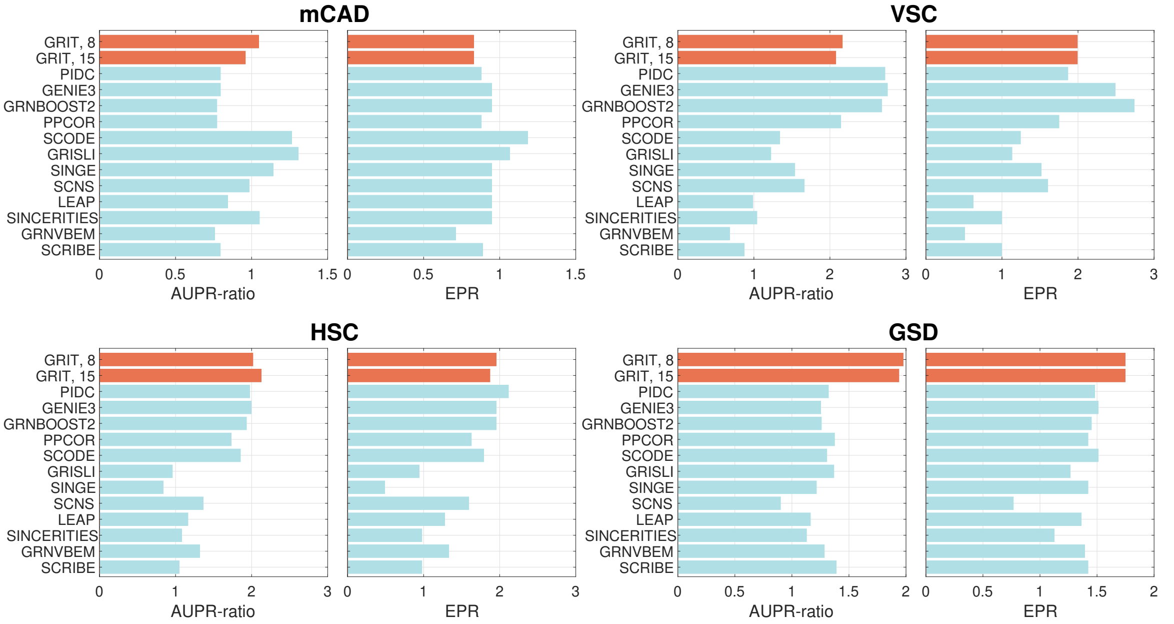


**Supplementary figure 3:** Detailed results of the BEELINE curated dataset. Bars show the median AUPR-ratios and EPRs for cases without dropouts (corresponding to [8, Fig. 4]). Results for the other methods are taken from the source file for [8, Fig. 4].

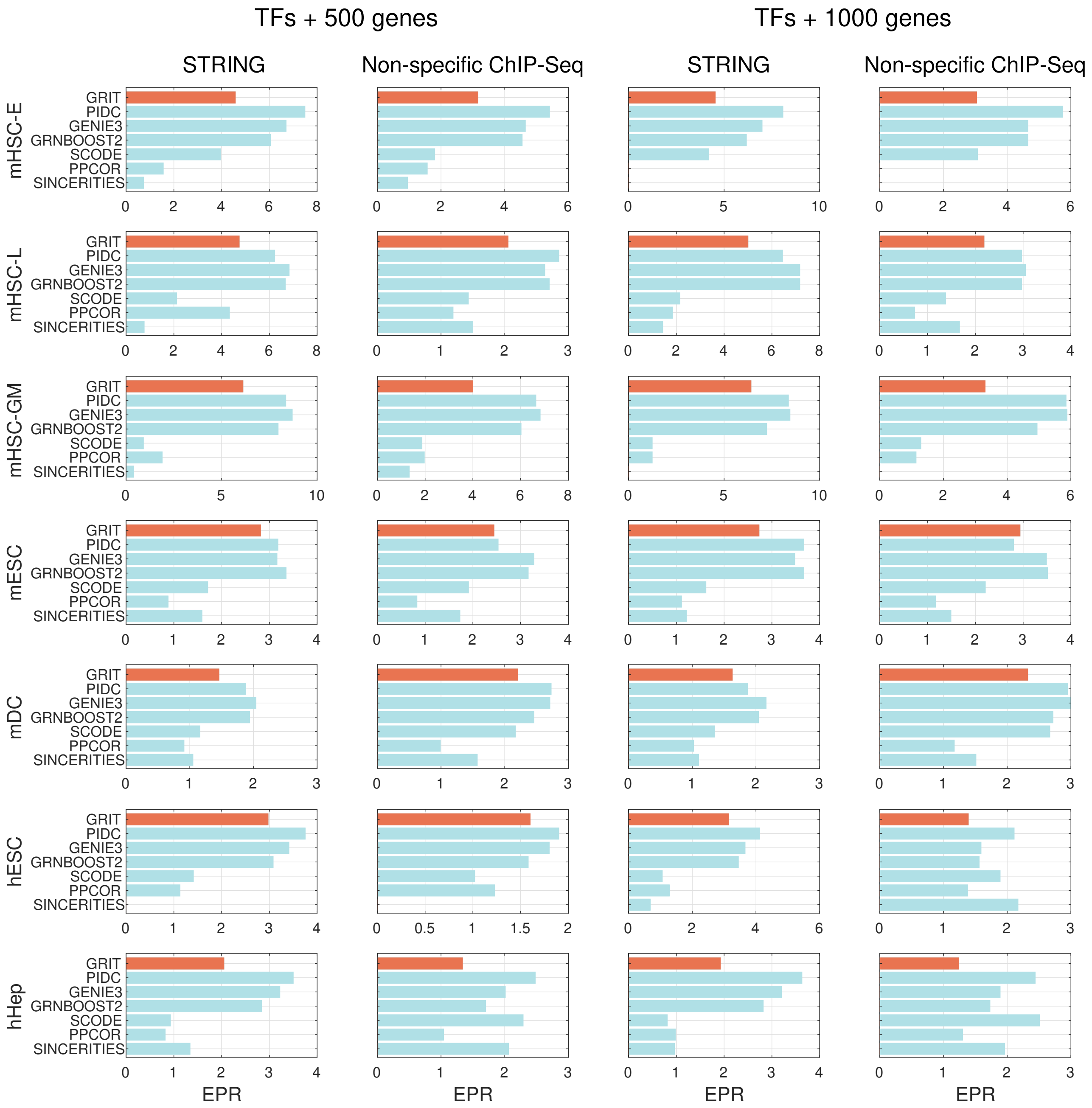


**Supplementary figure 4:** Detailed results of the BEELINE RNA-Seq datasets with the STRING and Non-specific ChIP-Seq ground truth networks (that seem to be the best ones for comparison). Complete results corresponding to [8, Fig. 5 and Suppl. Fig. 8] are in Supplementary table 2. Results for the other methods are taken from the source file for [8, Fig. 5].

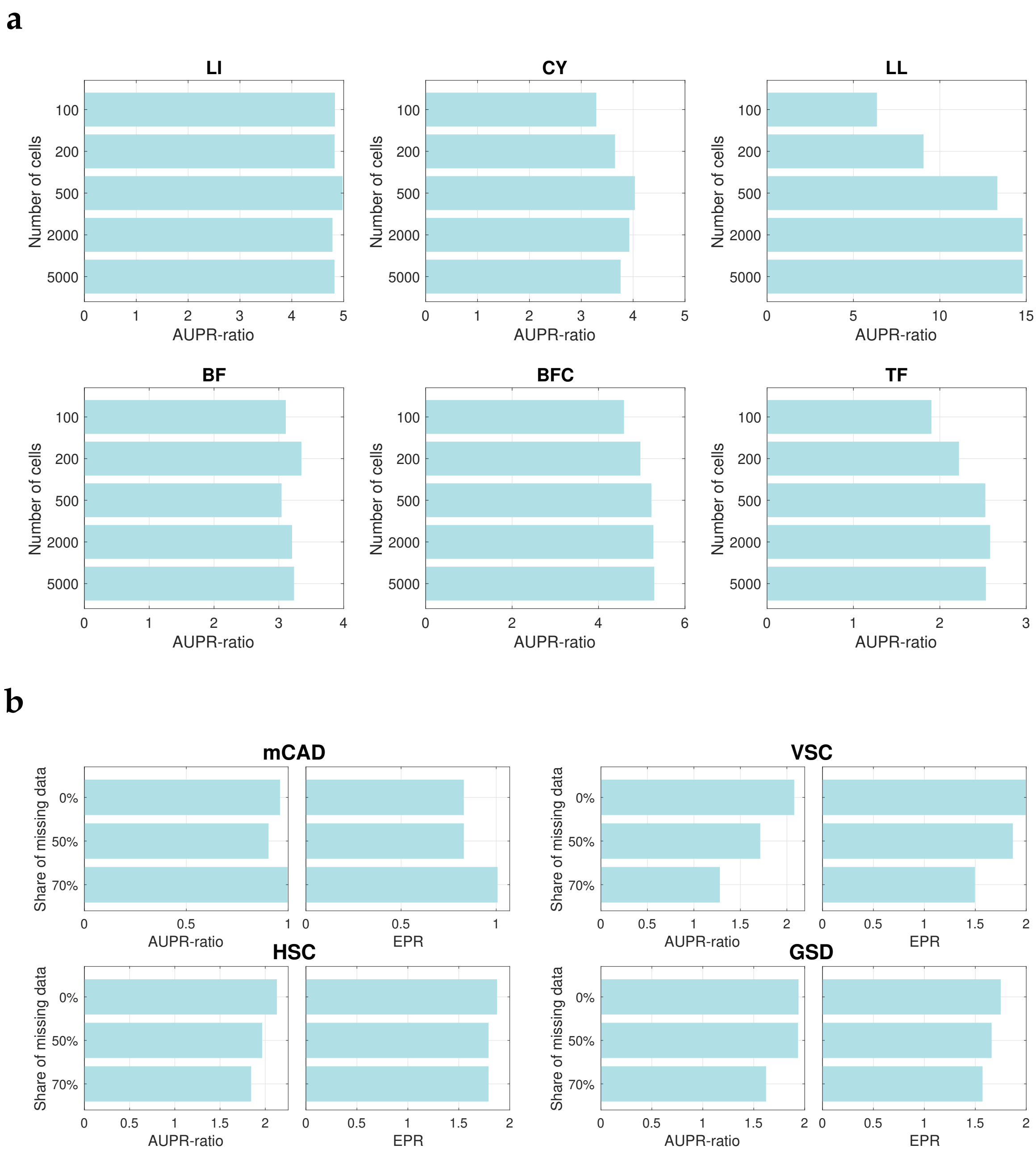


**Supplementary figure 5: a:** Effect of number of cells on the inference results of GRIT with 15 time points in the BEELINE synthetic dataset. **b:** Effect of dropout rate on the inference results of GRIT with 15 time points in the BEELINE curated dataset. The results are included for transparency, but it should be noted that the dropouts in BEELINE are generated by randomly selecting entries in the data matrices to be replaced by zero. This is not a realistic dropout model.

**a**

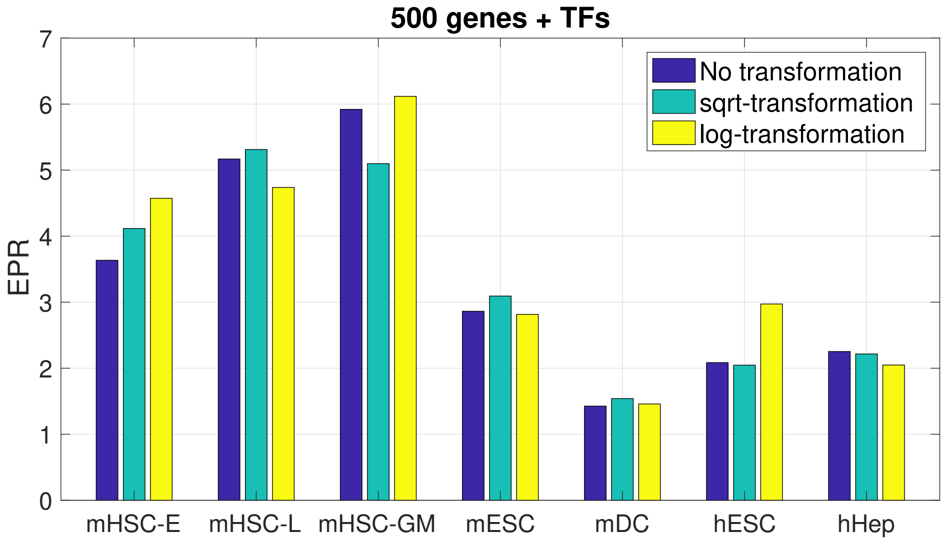


**b**

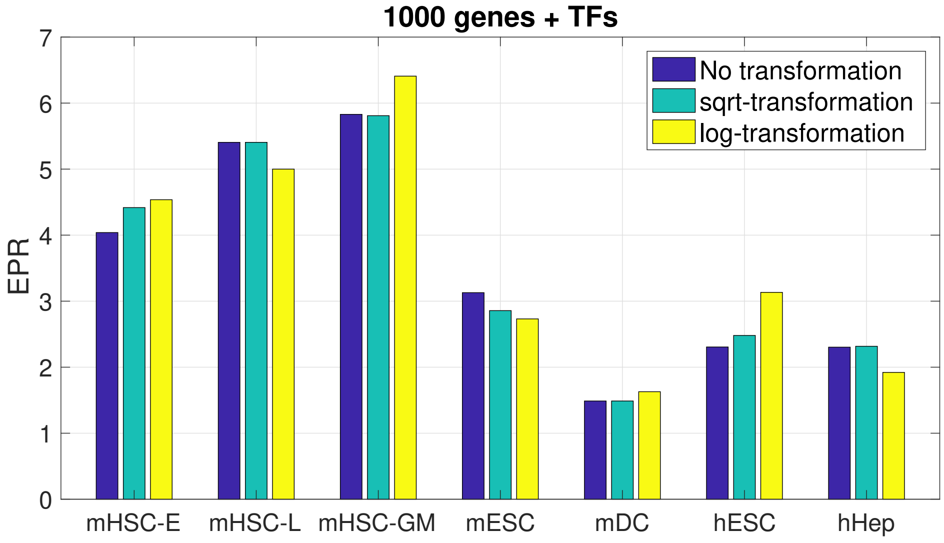


**Supplementary figure 6:** Effect of different transformations on the data on the inference results (EPR for STRING ground truth) for the case with TFs and 500 most highly varying genes (**a**) and TFs and 1000 most highly varying genes (**b**). The data as it is provided in BEELINE has been transformed by $y = \log_2(x+1)$. The case with no transformation is done by cancelling the log-transformation by calculating $x = 2^y - 1$ for each entry $y$ in the data matrix.

**a**

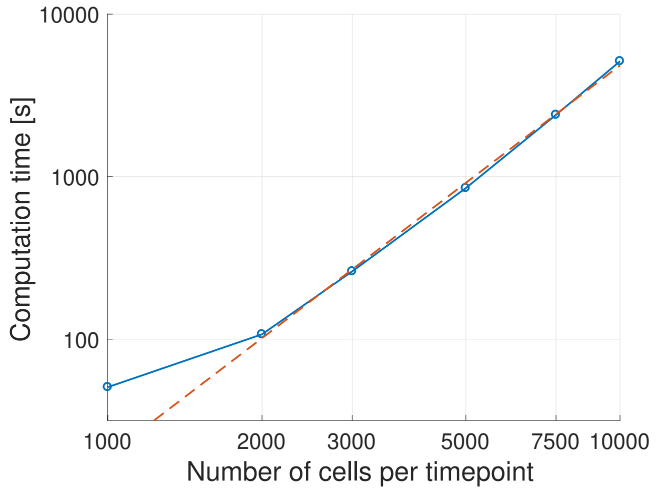


**b**

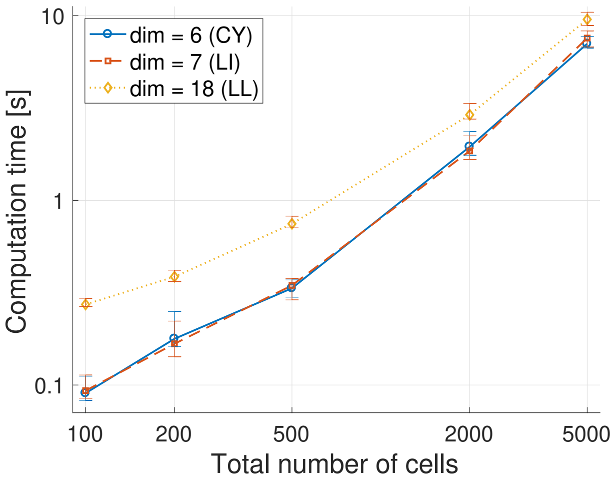


**Supplementary figure 7: a:** Computation times as a function of number of cells per timepoint in the experiment of Section 2.2 of the main text. The red dashed line is a regression line fitted to the cases with $\geq$ 2000 cells per timepoint. The regression line corresponds to $T \propto N_{\text{cell}}^{2.42}$ where $N_{\text{cell}}$ is the number of cells per timepoint. The times were very consistent between the five replicates and therefore only the means are shown in the plot. **b:** Computation times in the BEELINE synthetic dataset for the systems without branching dynamics. The computation time depends on the combination of system dimension and the number of cells. The case where the number of cells is very low (100), should be the most representative on the effect of dimension on computation time. In this case, the increase of the dimension from 6 or 7 to 18 increases the median computation time by a factor of 2.3 (0.1227$s$ or 0.1248$s$ to 0.2833$s$).
